# A population based expression atlas provides insights into disease resistance and other physiological traits in cassava (*Manihot esculenta* Crantz)

**DOI:** 10.1101/2021.09.23.461500

**Authors:** Alex C. Ogbonna, Punna Ramu, Esuma Williams, Leah Nandudu, Nicolas Morales, Adrian Powell, Robert Kawuki, Guillaume Bauchet, Jean-Luc Jannink, Lukas A. Mueller

## Abstract

Cassava, a food security crop in Africa, is grown throughout the tropics and subtropics. Although cassava can provide high productivity in suboptimal conditions, the yield in Africa is substantially lower than in other geographies. The yield gap is attributable to many challenges faced by cassava in Africa, including susceptibility to diseases and poor soil conditions. In this study, we carried out 3’RNA sequencing on 150 accessions from the National Crops Resources Research Institute, Ugandan for 5 tissue types, providing population-based transcriptomics resources to the research community in a web-based queryable cassava expression atlas. Differential expression and weighted gene co-expression network analysis were performed to detect 8,820 significantly differentially expressed genes (DEGs), revealing similarity in expression patterns between tissue types and the clustering of detected DEGs into 18 gene modules. As a confirmation of data quality, differential expression and pathway analysis targeting cassava mosaic disease (CMD) identified 27 genes observed in the plant-pathogen interaction pathway, several previously identified CMD resistance genes and two peroxidase family proteins different from the *CMD2* gene. Present research work represents a novel resource towards understanding complex traits at expression and molecular levels for the development of resistant and high-yielding cassava varieties, as exemplified with CMD.

## Introduction

Cassava (*Manihot esculenta* Crantz), a staple for over 800 million people worldwide, is cultivated across the tropics, with Africa accounting for over 50% of the total world production. Yield in Africa has remained substantially lower compared to other regions where cassava is grown^1^. Cassava has become a multipurpose crop with the ability to respond to the challenge of climate change and the potential to respond to priorities of developing countries including food security, poverty alleviation and economic development^1^. Most agronomic and production traits, such as yield, quality and disease-related traits, have become part of the primary breeding objective of a cassava breeding program and define the adoption of new cassava varieties by farmers and the market value of harvested roots^2,3^. The improvement of agronomic and production traits are enabled by improved understanding of the development and physiology characteristics of cassava, as reported for “leaf natural shading” using transcriptomics approach^4^. Transcriptomics is an approach that uses deep sequencing technologies such as the RNA-seq to profile transcriptomes, representing the complete set of transcripts in a cell^5^. Techniques such as transcriptomics can also be used to study plant diseases, such as Cassava mosaic disease (CMD). CMD, a major constraint to cassava production in Africa, Thailand and the Indian subcontinent^6,7^, with a yield loss of up to 95%, can be kept under control with the deployment of resistant varieties^8^. CMD is caused by several related species of cassava mosaic geminivirus (CGMs) and transmitted through infected cuttings and by a vector commonly known as whitefly (*Bemisia tabaci* G.). While much progress has been made on CMD, currently utilized resistance relies on single-gene resistance from related landraces. The narrow genetic base resistance could potentially break down over time, given their long-term effectiveness and the potential to be overcome by CGMs because of their fast-paced evolutionary rate^9,10^. On the other hand, the genetic mechanism of cassava brown streak disease (CBSD) - a threat to economic and food security for smallholder farmers in sub-Saharan Africa - is yet to be fully understood^11^, due to the difficulty in phenotyping the disease^12^ and the complex nature of CBSD virus resistance^13^. Recent efforts on identifying sources of resistance to CBSD using transcriptomics have been based on individual contrasting varieties^13,14^. Previous studies utilizing whole transcript sequencing technology to characterize and quantify transcripts have relied on comparison of transcriptomes allowing the identification of genes that are differentially expressed in response to individuals treated differently or individuals with contrasting characters of interest^15^. Fragmentation and library construction can introduce biases in whole transcript sequencing, bringing about more reads being assigned to a longer transcript than a shorter transcript, given that longer transcripts are sheared into more fragments^5^. The bias of fragmentation has been shown to enrich the differential expression of genes with longer transcripts^16^. 3’-RNA sequencing now provides a lower-cost and higher-throughput alternative to whole transcript sequencing, minimizing the aforementioned bias, and has been shown to have similar reproducibility and the ability to detect shorter transcripts^17^. This study is the first to use 3’-RNA sequencing technology in cassava, in a manner similar to earlier applications in maize^18^. For most complex traits, multiple small effect genetic variants can play a significant role in explaining trait variation when compared to simpler traits with rare monogenic mutations of large effects^19^. Therefore, analyzing gene expression levels of multiple tissues on a population basis would establish a high-resolution transcriptome resource for eQTL detection or trait prediction.

We present a population-based transcriptomic resource and expression atlas visualization for a population consisting of 150 cassava accessions sampled across five tissues (leaf, stem, fibrous root, storage root, flower) for studies of complex traits in the cassava community. The objectives of this study were to (i) quantify expression of transcripts across five tissues for 150 accessions, (ii) make this data resource available to the community in a web-based queryable cassava expression atlas, (iii) conduct differential gene expression analysis to detect differentially expressed genes (DEGs) across our population, with which we carried out weighted gene co-expression network analysis (WGCNA) and Gene Ontology (GO) analysis to characterize genes detected in different modules or co-expressed clusters, (iv) confirm data quality by differential gene expression and GO analysis carried out on clones differing for CMD tolerance. Altogether this work provides a population-based transcriptomics resource with a wide range of applications and can be leveraged for studies of simple and complex traits in cassava.

## Results

### Principal component analysis (PCA) highlights the clustering of different tissue types

RNA expressions from 150 accessions were quantified using the 3’-RNA sequencing method^20^. Five tissue types were profiled including storage root, fibrous root, stem, leaf, and flower for 150 cassava accessions, giving a total of 750 samples. PCA was performed to determine sample clustering. Using the variance stabilizing transformation (vst) normalized gene-level counts from HTSeq (Supplementary Table S1), PCA results indicated that samples of the same tissue types clustered together with PC1 and PC2 explaining 25.6 and 9.8% of the total variance in the gene expression across all tissues (Fig. 1a). For cassava accessions that did not flower at the time of sample collection or as a result of asynchronous flowering, tissue samples collected from their apical meristem clustered together with samples from flower tissues. Apical meristem tissue types were merged to flower tissue type for further downstream analysis. In agreement with PCA observations, heatmap and hierarchical clustering across the five tissue types demonstrated the same tissue type clustering and expression patterns (Fig. 1b).

**Figure 1.**
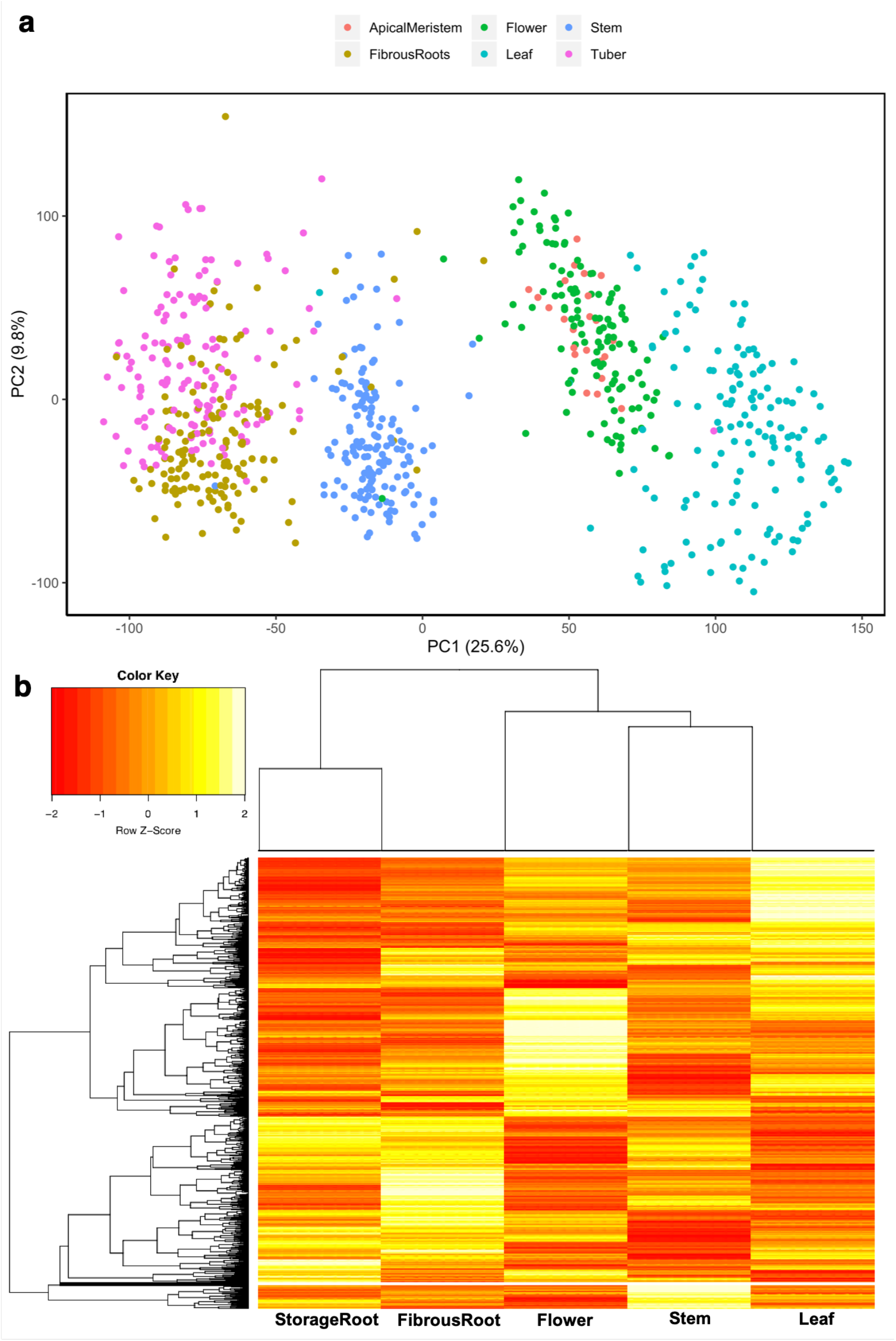
Principal component analysis and heatmap visualization of variance stabilizing transformation (vst) normalized gene-level counts for 31,895 genes across five different tissues (storage root, fibrous root, flower + apical meristem, stem and leaf). (A) Principal component 1 (PC1) and principal component 2 (PC2) were estimated using the prcomp function in R. The total variance explained by PC1 and PC2 is shown. (B) Heatmap of genes across different tissue types. Since apical meristem clustered with flower tissue type and was collected for accessions that did not flower, tissue samples from apical meristem were added to flower tissue types.

### Cassava expression atlas (CEA), a tool for visualizing quantified transcriptomes

The cassava expression atlas was implemented on cassavabase (https://cassavabase.org), an open-source digital ecosystem dedicated to the cassava research community^21,22^. Cassavabase provides a tool suite to assist breeders and breeding programs in automating their routine breeding and pre-breeding activities. Counts per million mapped reads (CPM) obtained from HTSeq gene-level counts normalized using EdgeR^23^ were used as digital units of expression for visualization of transcript expression levels (see Supplementary Table S2 for CPM values). Supplementary Fig. S1 provides a brief guide for using CEA functions. The CEA gene discovery search results include expression cube, expression images, heatmap and scatter plot features. To demonstrate CEA functions, we selected a random gene (Manes.01G011400.v6.1) and four accessions, and went through the process described in Supplementary Fig. S1 legend. The results from the different features are highlighted in Fig. 2. Figure 2 represents the expression cube, heatmap, images and barplot of the different tissue types for the gene of interest and other genes correlated to it, based on *r*^*2*^ correlation coefficient of 0.65 and above.

**Figure 2.**
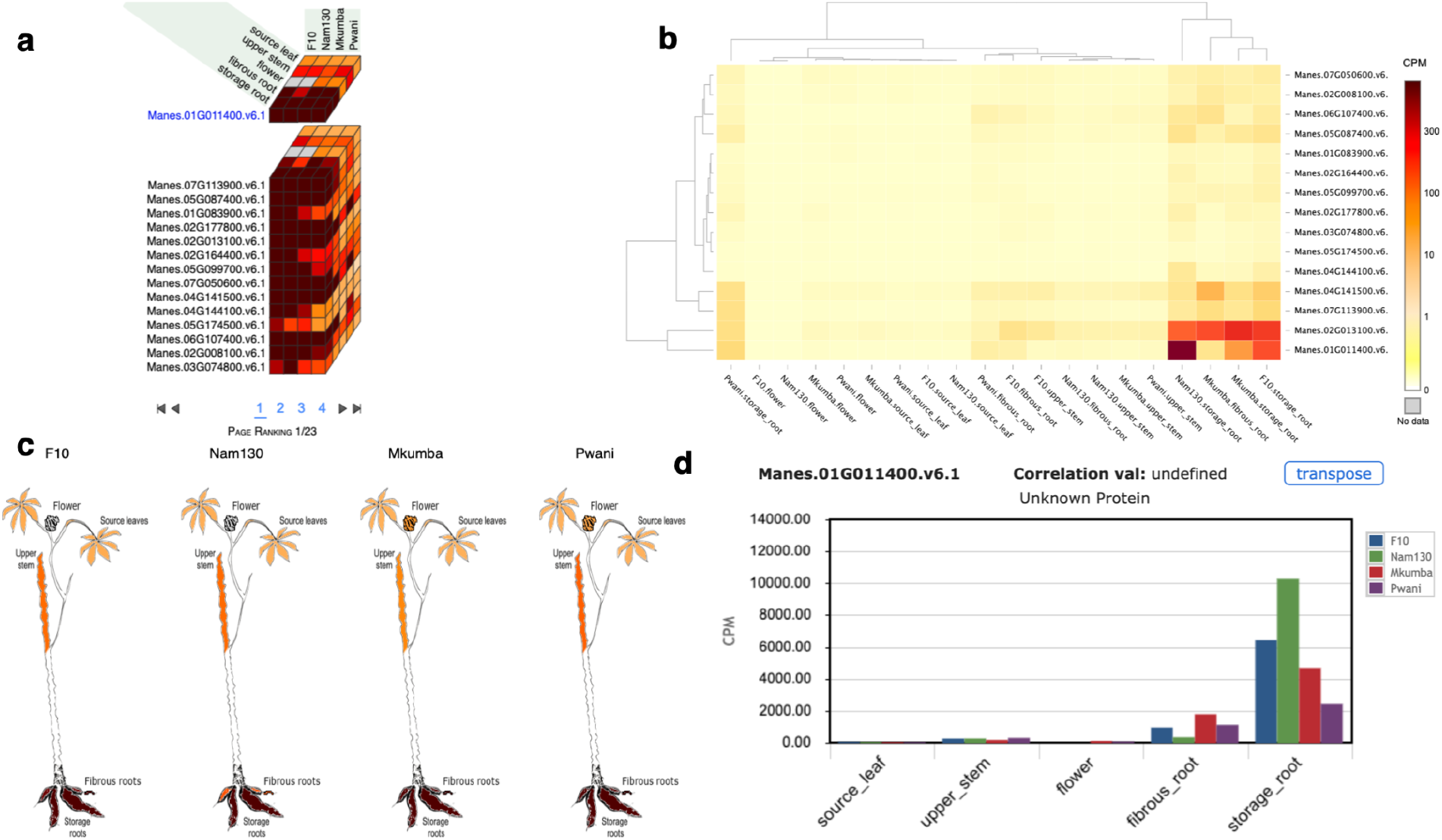
Cassava expression atlas (CEA) visualization output for four accessions (F10, Nam130, Mkumba, Pwani) and the Manes.01G011400 gene. (A) Expression atlas cube showing the expression of five tissue types on four accessions for Manes.01G011400 gene and other genes that are correlated to the gene of interest. (B) Heatmap of the expression of five tissue types on four accessions for Manes.01G011400 gene. (C) Expression images of five tissue types on four accessions for Manes.01G011400 gene. (D) Barplots showing the expression of five tissue types on four accessions for Manes.01G011400 gene.

### Differential expression analysis identifies genes involved in systemic acquired resistance

To quantitatively evaluate and compare transcript levels between tissue types, differential expression analysis was carried out using DESeq2 across the 150 accessions in our population. On average, 31,895 genes were mapped to at least one read in each of the tissue samples (Supplementary Table S3). A total of 19,445 DEGs were detected across pairwise tissue comparisons using an adjusted p-value of < *0.05* (Supplementary Fig. S2 and S3, Supplementary Table S4), yielding a maximum of 9,225 (storage root vs leaf, annotated as TvL) and a minimum of 3,330 (stem vs fibrous root) DEGs (Fig. 3a). However, for downstream analysis, a unique set of 8,820 unique DEGs were selected out of the 19,445 detected DEGs (Supplementary Table S5). The comparison of DEGs identified across contrasted tissue types show overlap of genes commonly expressed across contrasted groups, with larger numbers of genes overlapping across all contrasted groups for any specific tissue type (43.06% - 56.74%) (Fig. 3b-f). As a quality assessment positive control step, visualization of Rubisco methyltransferase family protein and Rubisco activase protein on CEA in a comparison between storage root versus leaf tissue types show that they are significantly differentially expressed (Supplementary Fig. S4). These genes are known to be housekeeping genes; Rubisco methyltransferase family protein and Rubisco activase protein^24^. Both proteins were significantly highly expressed in the leaf compared to other tissue types (Supplementary Fig. S4), as previously reported for *Arabidopsis thaliana^25^*.

**Figure 3.**
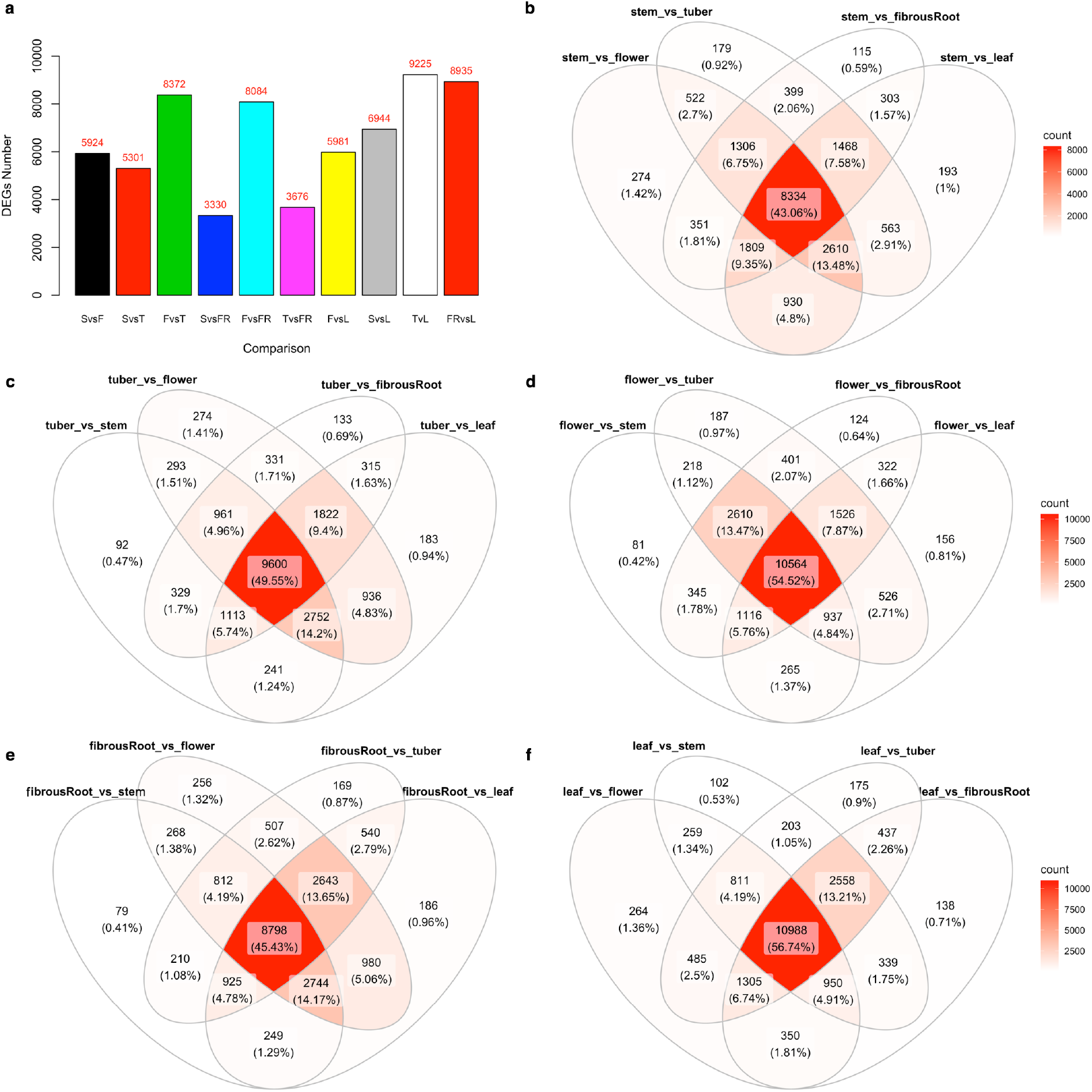
Detected differentially expressed genes (DEGs) across tissue types comparisons. (A) barplot showing detected DEGs across pairwise tissue types (SvsF = stem vs flower; SvsT = stem vs storageRoot; FvsT = flower vs fibrousRoot; TvsFR = storageRoot vs fibrousRoot; FvsL = flower vs leaf; SvsL = stem vs leaf; TvsL = storageRoot vs leaf; FRvsL = fibrousRoot vs leaf). (B-F) Overlaps of detected DEGs for different tissue type comparisons.

In order to confirm the high quality of our dataset, we carried out differential expression analysis between Mkumba and NASE14. Mkumba is a CMD resistant variety, while NASE14 expresses *CMD2*-mediated resistance as a mechanism for CMD resistance^26^. The implication of the *CMD2*-mediated resistance mechanism is that before the R-gene is activated, CMD viral infection leads to numerous molecular and physiological changes that can lead to limited symptom expression. Molecular screening indicated that resistant cultivars showed CMD symptoms at the early growth stages but the disease did not advance because of the presence of the *CMD2* resistance gene^27^. The samples used in this study were collected at early the growth stages with the aim of quantifying gene expressions, hence NASE14 was used as the “susceptible” variety. Since the effector triggered immunity was yet to be triggered but was rather at the phase of effector triggered susceptibility thus explaining the gene changes that typically occur in a susceptible clone^28^.

A total of 2,541 DEGs were detected between Mkumba and NASE14, based on adjusted p-value of < *0.05*, with 13% (321) and 87% (2,220) DEGs upregulated and downregulated, respectively (Fig. 4a, Supplementary Fig. S5, Supplementary Table S6). KEGG^29,30^ gene enrichment pathway analysis on the detected DEGs, identified genes (highlighted using red star) involved in the Plant-pathogen interaction pathway (27 DEGs; 1.6 fold enrichment) based on p-value of < 0.05 (Fig. 5, Supplementary Table S7). In the plant-pathogen interaction map, only two nodes (MKK1/2 and HSP90) were upregulated, the other 10 nodes were downregulated (Fig. 5a,c) based on log2 fold change. Hierarchical clustering of these identified plant-pathogen interaction pathway genes showed three clusters of genes based on their expression patterns across the gene clusters, with distinct expression across different tissue types (Fig. 5b). Other identified pathways included Pyruvate metabolism (15 DEGs), Base excision repair (9 DEGs) and Peroxime pathways with respective fold enrichments of 2.0, 2.4, and 1.8, and p-value of < 0.05. Based on Bonferroni multiple test correction of < 0.05, KEGG biological processes GO terms, showed that response to chitin was significantly enriched with 2.6 fold enrichment, and molecular function GO terms showed that protein binding, helicase activity, ATP binding, and metal ion binding were significantly enriched with fold enrichment of 1.4, 3.1, 1.3, and 1.3, respectively.

**Figure 4.**
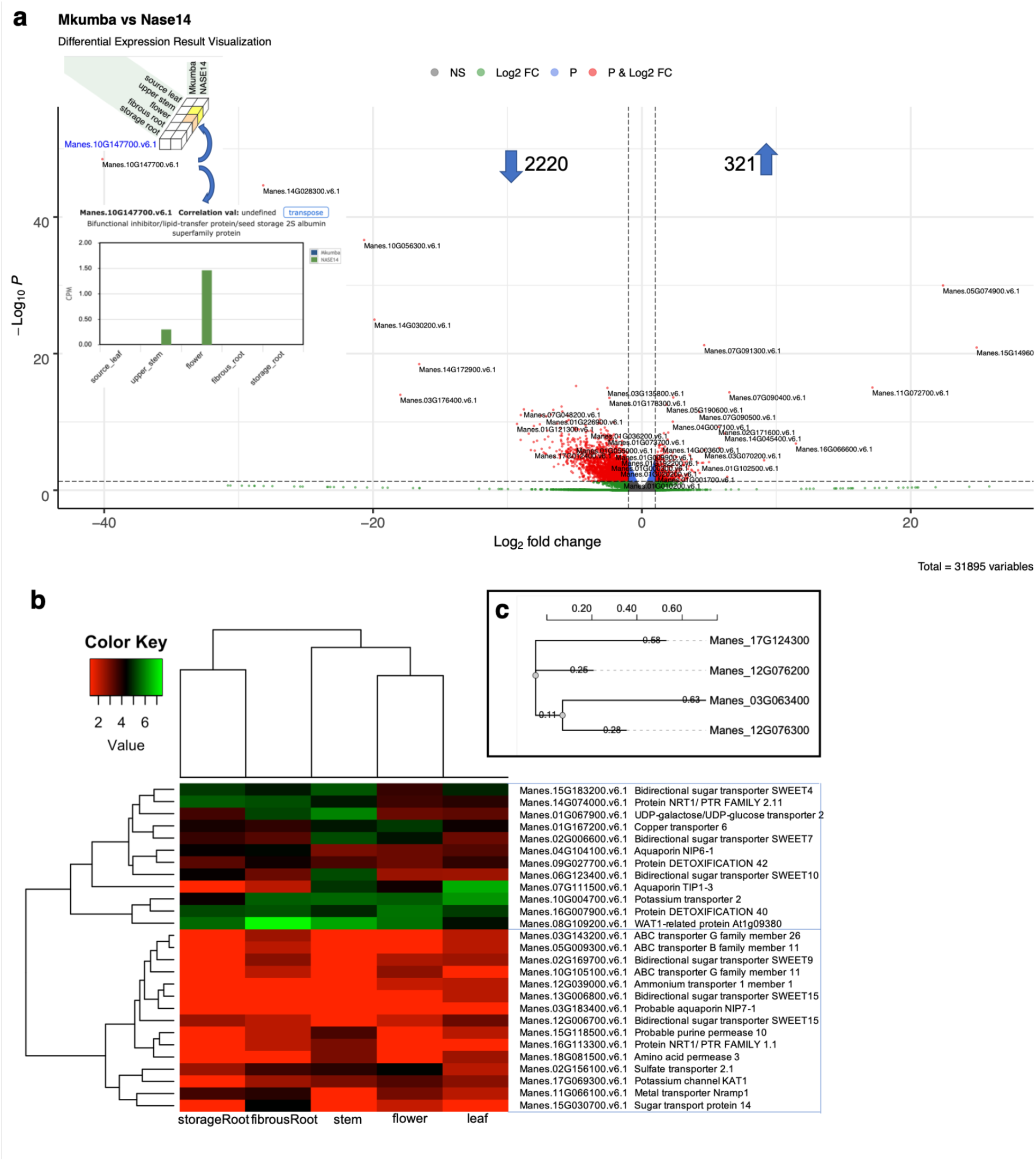
Differential expression between Mkumba vs Nase14, heatmap of hierarchical clustering and phylogenetics analysis of CMD associated peroxidase genes. (A) Volcano plot showing differentially expressed genes. Some of the gene names were printed on the plots. NS = not significant; Log2 FC = significant DEGs above the threshold of >|1| log2 fold change; P = significant DEGs based on adjusted p-value of < 0.05; P & Log2 FC = significant DEGs based on adjusted p-value of < 0.05 and threshold of >|1| log2 fold change. The gene (Manes.10G147700) in the blue ellipsis is a Bifunctional inhibitor/lipid-transfer protein/seed storage 2S albumin superfamily protein. Inserted are Expression atlas cube [above gene name] and barplot [below gene name] showing the expression of five tissue types on two accessions for Manes.10G147700 gene. (B) Heatmap for 27 genes involved in transmembrane transport activities in the yellow module with the genes and their corresponding description. The heatmap shows two broad characterizations. (C) Phylogeny of two differentially expressed peroxidase gene families and the two from GWAS analysis associated with CMD resistance in cassava (Wolfe et al. 2016).

**Figure 5.**
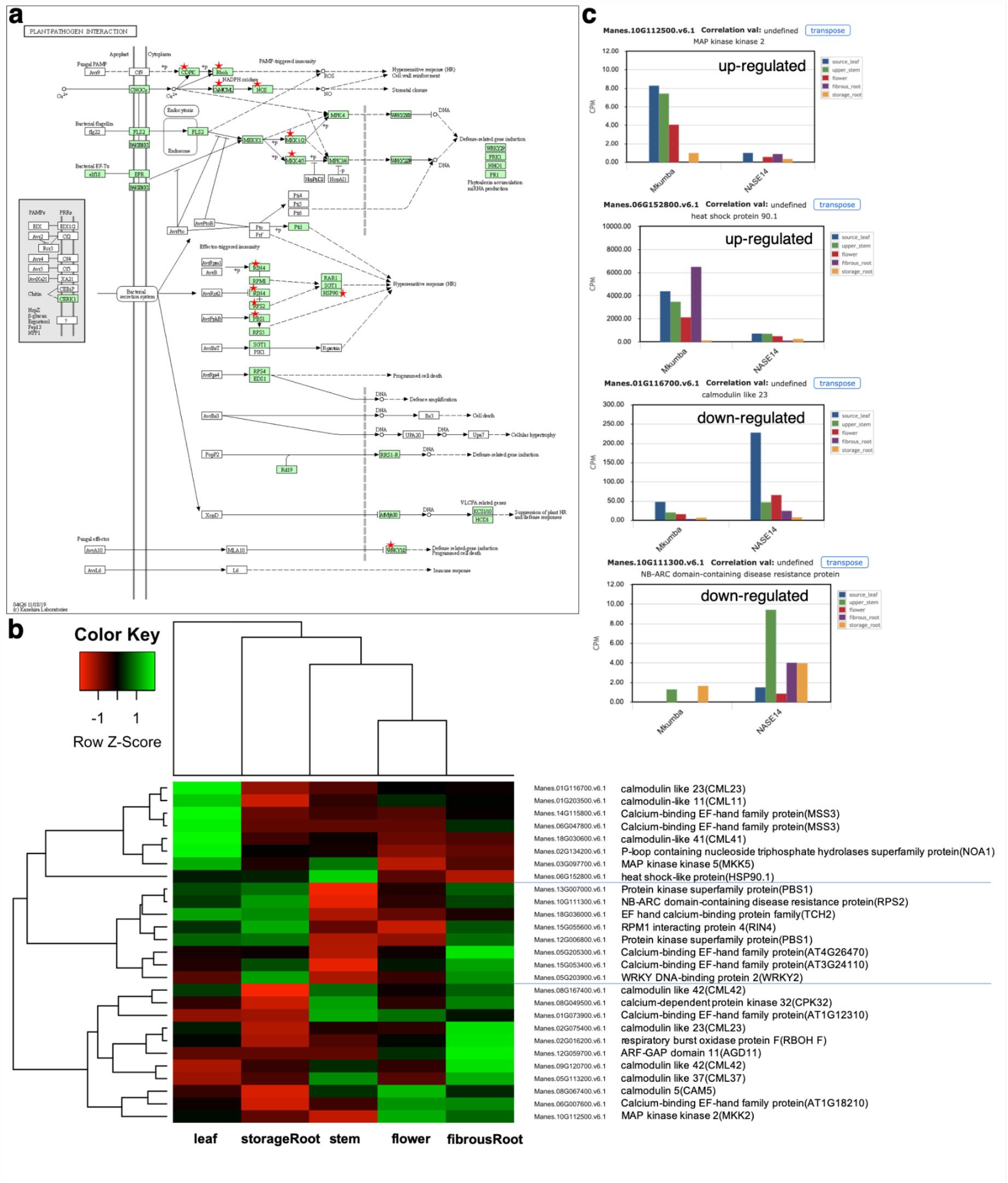
Plant-pathogen interaction pathway, Heatmap of hierarchical clustering of the 27 DEGs in the pathway and cassava expression atlas (CEA) barplot of selected genes. (A) Observed plant-pathogen interaction pathway revealed after the KEGG pathway enrichment analysis. Proteins present in the CMD DEGs are marked with red stars. Proteins marked in green belong to the reference-organism path. (B) Heatmap of hierarchical clustering for the 27 genes observed in the plant-pathogen interaction pathway using KEGG enrichment analysis. (C) barplot of some of the genes in the plant-pathogen interaction pathway showing the pattern of gene expression for genes that are upregulated and downregulated for CMD susceptible (based on *CMD2* resistance-NASE14) and resistant (Mkumba) accessions. The plant-pathogen interaction pathway reveals sets of genes involved in the plant immune response with MKK1/2 and HSP90 upregulated based on log2 fold change and showing down- and up-regulation in CMD susceptible and resistant accessions, respectively. The rest of the observed genes in the plant-pathogen interaction pathway were downregulated based on log2 fold change. The functions of these genes included potential calcium sensors (AGD11, CML, CAM), the second phase of hydrogen peroxide generation during hypersensitive response-like cell death (MKK5), and disease resistance pathogen recognition protein that triggers a defense system including the hypersensitive response, which restricts the pathogen growth (RPS2)^84^. Others were genes that produce nitric oxide (NO) which is a messenger molecule involved in hormonal signaling and defense responses in plants (NOA1), a protein kinase gene involved in plant defense responses specifically recognizing effector avirulence protein and triggering a defense reaction (RPS5), and a gene that is an essential regulator of plant defense, which plays a central role in resistance in case of infection by a pathogen (RIN4). Additional genes in the pathway were involved in transcription that interacts specifically with the W box (5’-(T)TGAC[CT]-3’) (WRKY2), and a gene that generates reactive oxygen species (ROS) during incompatible interactions with pathogens and is important in the regulation of the hypersensitive response (HR) (RBOH F). The observed enriched GO terms were biological processes that were part of the plant-pathogen interaction pathway.

### Weighted gene co-expression network analysis (WGCNA) of the detected 8,820 unique DEGs identified 18 gene modules

WGCNA was carried out using the 8,820 DEGs obtained from comparison of different tissue types for construction of a scale-free co-expression network. These 8,820 DEGs showed distinct expressions across different tissue types (Fig. 6a). The weighted coefficient parameter **β** = 7 was chosen to obtain a correlation coefficient of 0.93 (Supplementary Fig. S6). A hierarchical tree function was used to divide the constructed clustered tree, detecting 18 co-expression modules, each tagged with a color, including turquoise with maximum number of 2,848 DEGs, light green with minimum number of 39 DEGs and a yellow module with 307 DEGs (see all modules in Fig. 6b-d, Supplementary Fig. S7a & Table S8).

**Figure 6.**
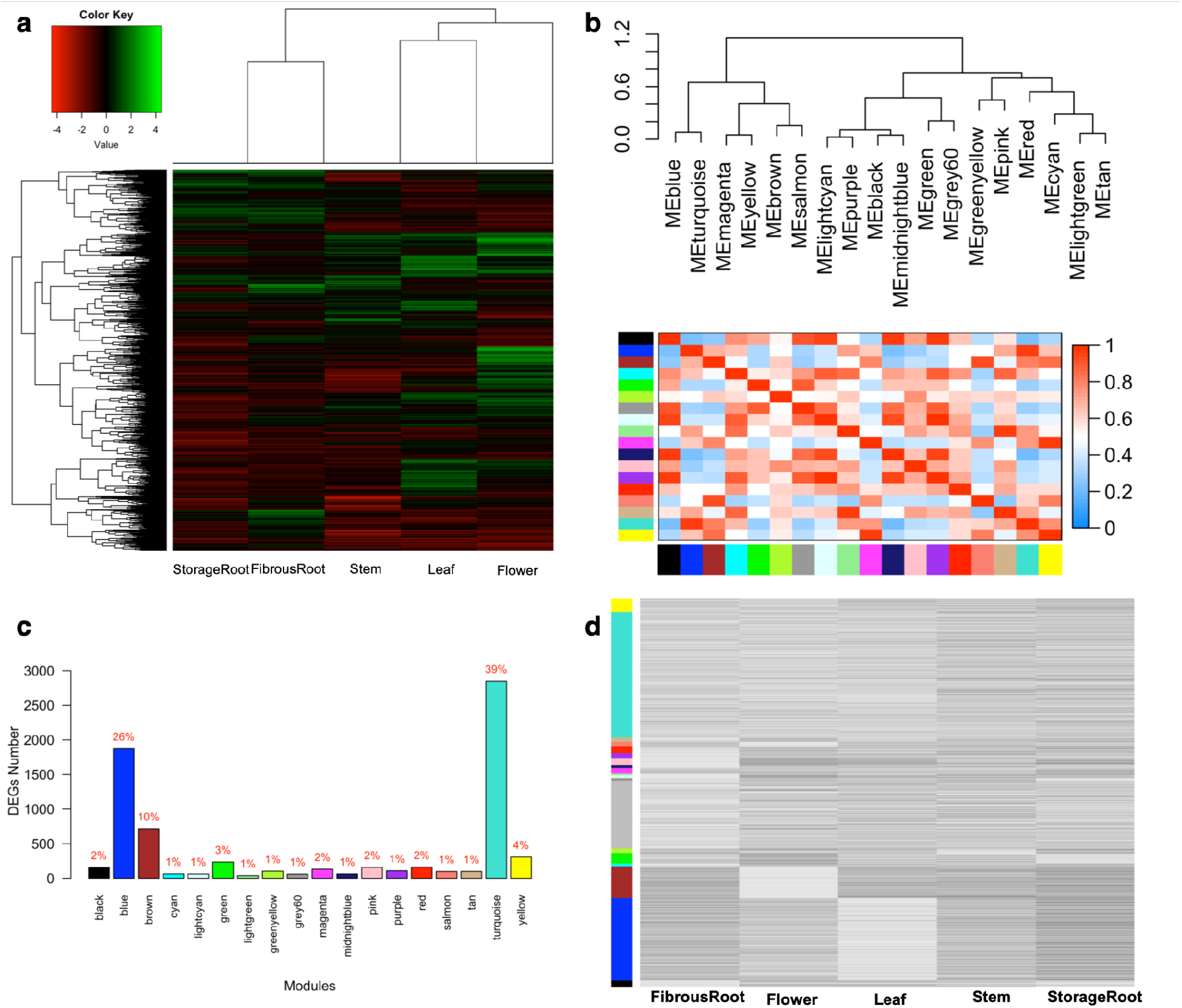
Heatmap of hierarchical clustering analysis and weighted gene co-expression network analysis (WGCNA). (A) Heatmap of 8,820 DEGs across different tissue types. (B) Gene co-expression modules showing the cluster dendrogram constructed based on the eigengenes of the modules (upper panel) and the heatmap for the correlation coefficient between the modules (lower panel). (C) Barplot showing the approximate percentage distributions of DEGs clustered into 18 gene modules using the WGCNA R package. (D) Expression patterns of genes as they are clustered based on detected modules and different tissue types. Genes in the grey module are usually genes that did not cluster with genes in any of the 18 modules.

Given that the yellow module with 307 DEGs contains a gene previously characterized as an intra-cellular transporter in a recent study conducted by our group, we subjected it to further analysis. First, we functionally classified the genes in the yellow module based on (1) molecular function, (2) biological function, (3) cellular components, (4) protein classes and (5) pathway categories using PANTHER classification system version 16.0^31^ with corresponding *Arabidopsis thaliana* annotation IDs from the cassava version 6.1 annotation genome^32^.

The molecular function categories observed for genes in the yellow module included Binding (GO:0005488), Catalytic activity (GO:0003824), Molecular function regulator (GO:0098772) and Transporter activity (GO:0005215) with 23%, 58.4%, 7.1% and 11.5% genes in each category, respectively (Supplementary Fig. S7b). The categories of protein classes represented in the yellow module included genes encoding metabolite interconversion enzymes (PC00262), transporters (PC00227) and gene-specific transcriptional regulators (PC00264) with 63%, 10.6%, and 8.55% genes in each category, respectively (Supplementary Fig. S8a). The biological process GO term categories for genes in the yellow module included cellular processes (GO:0009987), metabolic processes (GO:0008152) and biological regulations (GO:0065007) with 34%, 30.5% and 10.6% genes in each category, respectively (Supplementary Fig. S8b). The cellular component GO terms categories observed in the yellow module included cellular anatomical entities (GO:0110165) and intercellular activities (GO:0005622) with 58.3% and 39.6% of genes for each category, respectively (Supplementary Fig. S8c). Pathway characterization shows that genes in the yellow module fall within categories including glycolysis (P00024), fructose galactose metabolism (P02744), pentose phosphate pathway (P02762), and vitamin B6 metabolism (P02787) with gene percentage of 21.1%, 10.5%, 10.5%, 10.5%, and 5.3%, respectively (Supplementary Fig. S8d). Other pathway categories included cholesterol (P00014), cysteine (P02737), serine glycine (P02776), and tyrosine (P02784) biosynthesis, all at 5.3% of genes in the yellow module (Supplementary Fig. S8d).

Second, the yellow module analysis for enriched GO terms was conducted for (1) biological processes, (2) molecular functions and (3) cellular components based on the Bonferroni-corrected p-value of < 0.05. The statistically significant enriched terms indicated that genes in the yellow module were mostly involved in biological processes of sporopollenin biosynthesis (p-value = 2.69E-06), anther wall tapetum development (p-value = 0.0065), cuticle development (p-value = 0.0045), cellular carbohydrate catabolic process (p-value = 0.00086), and starch metabolic process (p-value = 0.0064; Table 1) (with all GO lists in Supplementary Table 9). For molecular function GO terms, genes in the yellow module were mostly involved in sugar transmembrane transporter activity (p-value = 0.0036), oxidoreductase activity (p-value = 0.01), iron ion binding and heme binding (p-value = 0.01). For cellular component enriched GO terms, integral component of plasma membrane (p-value = 0.033), plant-type cell wall (p-value = 0.0006) and protein-containing complex (p-value = 0.0007) were statistically significantly enriched terms for genes in the yellow module (Supplementary Table S9).

**Table 1.** List of six genes from the yellow module identified in the biological processes GO to be involved in starch metabolic processes using PANTHER overrepresentation Fisher’s exact test with Bonferroni correction for multiple testing for p-value < 0.05.

| No | GeneID | Annotation | Gene name |
| --- | --- | --- | --- |
| 1 | Manes.14G031100.v6.1 | AT5G51820 | phosphoglucomutase |
| 2 | Manes.01G055700.v6.1 | AT1G32900 | UDP-Glycosyltransferase superfamily protein |
| 3 | Manes.01G236700.v6.1 | AT1G27680 | ADPGLC-PPase large subunit |
| 4 | Manes.06G021000.v6.1 | AT3G52180 | dual specificity protein phosphatase (DsPTP1) family protein |
| 5 | Manes.18G063500.v6.1 | AT4G09020 | isoamylase |
| 6 | Manes.03G171500.v6.1 | AT3G55760 | Uncharacterized protein |

A set of 27 genes from the yellow module were found to be involved in transmembrane transporter activities in the molecular function GO using PANTHER gene enrichment tool with Fisher’s exact test and Bonferroni correction for multiple testing for p-value of < 0.05, included Manes.16G007900, a MATE transporter, earlier reported to be involved in the regulation of cyanogenic glucosides in cassava root^33^ (Table 2). The expression pattern of these sets of 27 genes shows at least two broad groups with genes mostly involved in detoxification found in the upper half (Fig. 4b). The GO term enrichment results for all detected modules are described in Supplementary Table S10.

**Table 2.** List of 27 genes involved in transmembrane transporter activity identified in the molecular function GO for yellow module genes, using PANTHER overrepresentation Fisher’s exact test with Bonferroni correction for multiple testing for p-value < 0.05. Manes.16G007900 MATE transporter, earlier reported to be involved in the regulation of cyanogenic glucosides in cassava root, was found in these groups of genes.

| No | GeneID | Annotation | Gene name | Protein class |
| --- | --- | --- | --- | --- |
| 1 | Manes.05G009300.v6.1 | AT1G02520 | ABC transporter B family member 11 | ATP-binding cassette (ABC) transporter |
| 2 | Manes.08G109200.v6.1 | AT1G09380 | WAT1-related protein At1g09380 |  |
| 3 | Manes.10G105100.v6.1 | AT1G17840 | ABC transporter G family member 11 | ATP-binding cassette (ABC) transporter |
| 4 | Manes.09G027700.v6.1 | AT1G51340 | Protein DETOXIFICATION 42 | transporter |
| 5 | Manes.15G030700.v6.1 | AT1G77210 | Sugar transport protein 14 | secondary carrier transporter |
| 6 | Manes.18G081500.v6.1 | AT1G77380 | Amino acid permease 3 |  |
| 7 | Manes.04G104100.v6.1 | AT1G80760 | Aquaporin NIP6-1 |  |
| 8 | Manes.11G066100.v6.1 | AT1G80830 | Metal transporter Nramp1 | secondary carrier transporter |
| 9 | Manes.01G167200.v6.1 | AT2G26975 | Copper transporter 6 | secondary carrier transporter |
| 10 | Manes.02G169700.v6.1 | AT2G39060 | Bidirectional sugar transporter SWEET9 |  |
| 11 | Manes.10G004700.v6.1 | AT2G40540 | Potassium transporter 2 | transporter |
| 12 | Manes.03G183400.v6.1 | AT3G06100 | Probable aquaporin NIP7-1 |  |
| 13 | Manes.03G143200.v6.1 | AT3G13220 | ABC transporter G family member 26 | ATP-binding cassette (ABC) transporter |
| 14 | Manes.16G113300.v6.1 | AT3G16180 | Protein NRT1/ PTR FAMILY 1.1 | transporter |
| 15 | Manes.16G007900.v6.1 | AT3G21690 | Protein DETOXIFICATION 40 | transporter |
| 16 | Manes.15G183200.v6.1 | AT3G28007 | Bidirectional sugar transporter SWEET4 |  |
| 17 | Manes.07G111500.v6.1 | AT4G01470 | Aquaporin TIP1-3 |  |
| 18 | Manes.02G006600.v6.1 | AT4G10850 | Bidirectional sugar transporter SWEET7 |  |
| 19 | Manes.12G039000.v6.1 | AT4G13510 | Ammonium transporter 1 member 1 | primary active transporter |
| 20 | Manes.15G118500.v6.1 | AT4G18210 | Probable purine permease 10 |  |
| 21 | Manes.01G067900.v6.1 | AT4G23010 | UDP-galactose/UDP-glucose transporter2 | secondary carrier transporter |
| 22 | Manes.02G156100.v6.1 | AT5G10180 | Sulfate transporter 2.1 | transporter |
| 23 | Manes.12G006700.v6.1 | AT5G13170 | Bidirectional sugar transporter SWEET15 |  |
| 24 | Manes.13G006800.v6.1 | AT5G13170 | Bidirectional sugar transporter SWEET15 |  |
| 25 | Manes.17G069300.v6.1 | AT5G46240 | Potassium channel KAT1 | ion channel |
| 26 | Manes.06G123400.v6.1 | AT5G50790 | Bidirectional sugar transporter SWEET10 |  |
| 27 | Manes.14G074000.v6.1 | AT5G62680 | Protein NRT1/ PTR FAMILY 2.11 | transporter |

### Genetic variance explained by SNP markers in the detected 8,820 DEG regions

Evaluating the functional relevance of the detected DEGs, proportion of variance explained by SNPs in the DEG regions was higher for vigor (DEG SNPs: 0.24, sampled SNPs: 0.12), a fitness trait and dry matter (DM) content (DEG SNPs: 0.38, sampled SNPs: 0.32), an agronomic trait when compared to randomly selected SNPs (excluding SNPs in high LD with the DEG SNPs) of equal size and distribution (Supplementary Fig. S10). For other agronomic and disease related traits, SNPs within the DEG regions explained less variance when compared to the randomly sampled SNPs of equal sizes and distribution. While fitness related traits respond to gene regulation (though fresh root yield does not respond in that manner based on this dataset), disease resistance in plants has been shown not to be influenced by differential expression but rather by genes that are involved in disease infection recognition^18,34,35^. This is also highlighted by the SNPs in the DEG regions explaining about twice the variance explained by the rest of the genome (11,399 DEG-SNPs: 0.23; 65,054 Other-SNPs: 0.13) for plant vigor trait.

## Discussion

The multifaceted ability of cassava to respond to both the challenge of climate change and the priorities of developing countries makes cassava a sustainable and reliable crop for food security, poverty alleviation and economic development^1^. However, the cassava yield gap in the developing countries compared to other regions where cassava is grown highlights the necessity for better understanding of the cassava developmental and physiological processes. This will allow further improvements in complex traits including yield, quality, fitness and disease resistance-related traits in Sub-Saharan Africa. Here, we aimed to provide a population-based transcriptomics resource for studies of complex traits to assist cassava improvement efforts.

Expression of transcripts for 150 accessions from the Ugandan cassava breeding program were quantified, with sampling performed on flower, leaf, stem, fibrous and storage root tissues for each accession. A 3’-RNA sequencing method was chosen, which has been shown to be more efficient than RNA sequencing in the way it handles paralogs^20^. Multivariate analysis shows the clustering of each tissue type together. PCA outputs split variance according to distinct biological parameters, with PC1 highlighting the variance due to different tissue types, while PC2 likely explains the genetic differences between the accessions in the population. Similar observations were made using a heatmap hierarchical clustering algorithm, supporting our earlier PCA observation and highlighting similarities in the pattern of expression between tissue types, with the storage root having a closer expression pattern to the fibrous root, while the leaf had a similar expression profile to the stem tissue. The resolution of the different tissue types into distinct clusters indicated the high quality of the dataset, and highlights that the quantified transcripts captured transcriptional differences in the tissue types and the physical relatedness between tissue types. A similar clustering pattern was previously reported for cassava among tissue types^15^.

Differential expression results and overlapping of identified DEGs between contrasting tissue type comparisons, highlight common DEGs across comparisons and indicate the abundance of house-keeping genes and smaller percentage of genes that are unique or specific to a set of contrasted tissue types. In addition, it was observed that tissue types with similar expression patterns had fewer DEGs than those having more distinct expression patterns. For example, stem vs fibrous root tissue comparison gave the least number of DEGs (3,330), followed by storage root vs fibrous root comparison with 3,676 DEGs, while comparison of storage root vs leaf tissue gave the highest number of DEGs (9,225). This observation is consistent, supports our PCA and hierarchical clustering results, and represents the first study in cassava that looked at the comparative expression patterns of major organs/tissues using a population-based approach. In addition, expression patterns of the detected modules by tissue types seem to suggest modules/genes that could be targeted for cassava improvements based on traits and contrasting tissue types of interest. This highlights how genes within a module are differentially expressed across different tissue types. For example, genes in the brown module show that flower tissues had distinct expression patterns compared to the rest of the tissue types, while genes in the blue module show that leaf tissues had distinct expression patterns compared to the rest of the tissue types.

The classification of the genes in the yellow module indicates that they are mostly involved in regulatory functions such as RNA-& DNA-binding transcription and other transcription factors required for the regulation of many cellular processes, including transcription, translation, gene silencing, gene expression control, catalysis and similar functions^36^. Other genes in the yellow module are involved in transport activities, allowing the transfer of substances (sugar, copper, sulfate, potassium, iron) across plasma membranes, enabling activities such as detoxification as earlier described for cyanogenic glucosides in cassava^33^.

GO term enrichment (over-representation) analysis showed that genes in the yellow module are involved in both molecular, cellular and biological functional processes. Notable among them are the sugar transmembrane transporter activity, integral component of plasma membrane and sporopollenin biosynthetic process with the most upregulated fold enrichment, supporting the earlier speculation that most genes in this module are mostly associated with plant developmental processes. While Manes.10G147700 found in the yellow module, involved in plant defense against cassava mosaic virus and its insect vector whitefly, is characterized to be involved in lipid transport/protein metabolism based on functional GO terms. The cassava root HCN regulation gene (Manes.16G007900) also found in the yellow module, and part of the transmembrane transport activity GO term, supports the speculation that genes in the yellow module are functionally involved in developmental processes. Manes.16G007900 is a MATE transporter that may be involved in the regulation of cyanogenic glucosides such as linamarin and lotaustralin in cassava root^33^. Linamarin, an abundant cyanogenic glucoside variant in cassava and a secondary metabolite, contains nitrogen and serves as a nitrogen shuttling and storage compound^37^. Nitrogen is a vital and major component of chlorophyll used as an energy source to produce sugar, a major component of amino acids, the building blocks of proteins, and together they ensure the survival and development of a plant^38^.

Twenty-seven transporter genes from the yellow module, characterized to be involved in transmembrane transporter activity based on molecular function GO terms, facilitates transfer of a specific substance or group of related substances, from one side of a membrane to the other^39^. The expression pattern of these 27 transmembrane transporters indicated two broad categories: those likely involved in allowing the uptake of essential nutrients and those involved in secretion of metabolic end products & hazardous substances. Among those already characterized in cassava are Manes.15G030700 involved in galactose transport^40^ and Manes.16G007900 involved in cyanogenic glucoside transport^33^. However, the phosphoserine aminotransferase family gene (Manes.02G112800 - phosphoserine transaminase), identified to be involved in vitamin B6 metabolism, has not been previously characterized in cassava. Six genes in the yellow module characterized by PANTHER biological process GO appear to be involved in the starch metabolic processes, and two were previously characterized as being involved in the starch biosynthetic pathway in cassava^41,42^.

Differential expression results were further investigated for CMD tolerance. DEGs detected encoding NBS (Nucleotide Binding Site) and TIR (Toll/Interleukin-1 Receptor) regions were previously described by Lopez et al. as resistance gene candidates in cassava^43^. Eleven (11% R genes) out of the 99 resistance analogues genes reported by Allie et al. to respond to cassava mosaic virus infection were among the detected DEGs^44^ (Supplementary Table S11). The eleven R genes included leucine-rich repeat (LRR) transmembrane protein kinase and NB-ARC domain-containing disease resistance protein, disease resistance protein (TIR-NBS-LRR class) family and Histone superfamily protein. Previous genome-wide association studies (GWAS) on CMD resistance in African cassava identified two closely linked genes on chromosome 12, Manes.12G076200 and Manes.12G076300, both identified as peroxidase superfamily proteins to be involved in the regulation of CMD^9,10^. One is characterized as *CMD2* and the second is a “highly correlated peroxidase gene about 17 kilobases away”^45–49^. However, neither of these peroxidase genes were differentially expressed in our dataset, despite the fact that they were characterized as pathogenesis-related proteins involved in host response to infection^50^ and downregulated in response to CMD infection in susceptible accessions^44^. Surprisingly, we found two different peroxidase superfamily proteins, Manes.03G063400 (log2 fold change: −2.7919) and Manes.17G124300 (log2 fold change: 9.0907), located on chromosomes 3 and 17, respectively to be significantly differentially expressed. The percentage sequence similarity of the *CMD2* gene to Manes.03G063400 and Manes.17G124300 genes was 52.80% and 52.73%, and that of Manes.12G076300 to Manes.03G063400 and Manes.17G124300 genes were 53.06% and 48.12%, respectively. With an average of 50% pairwise sequence similarity across homologs (Fig. 4c), the significance of these findings will require additional investigation to ascertain if these genes are involved in CMD response.

Coincidentally, based on an optimal adjusted p-value (7.45E-45) and log2 fold change (−40.13), we identified a lipid-transfer protein (Manes.10G147700), previously reported to be involved in plant defense against cassava mosaic virus and its insect vector whitefly^8^. Vidya et al., used the Nucleotide Binding Site transcriptome profiling technique with the aim of identifying CMD resistance genes in India^8^. A total of 24 genes (27 DEGs with homologs) out of the 105 candidates genes reported to be associated with CMD resistance by Wolfe et al. were part of our DEGs, including Manes.10G147700 reported to be a candidate gene associated with CMD resistance when phenotyped at three months after planting (CMD3S)^10^ (Supplementary Table S11). These 24 genes reported by Wolfe et al., are associated with CMD resistance across all phenotyping time points (CMD1S, CMD3S, CMD6S, CMD9S)^10^. Manes.10G147700 was expressed in flower tissue and at a lower level in the stem of a susceptible cassava variety (see Fig. 4a inserted CEA cube and barplot), similar to the expression pattern observed in its Arabidopsis homolog (AT3G52130)^51^. Manes.10G147700 is a non-specific lipid transfer protein (nsLTP) with the ability to transfer lipids across membranes^52^. nsLTP has been previously described in *Arabidopsis*, maize, spinach, castor bean, wheat^53^ and only recently reported to be involved in CMD resistance in cassava^8^. A non-specific lipid transfer protein, encoded by large gene families in many flowering plants^54^, binds to sterol molecules to trigger plant defense response by interacting with a receptor at the plant plasma membrane^55^ and usually detected during early development in plants^56^. This protein was previously implicated in plant defense against viral, fungal and bacterial pathogens in plants^57^. Other identified DEGs included heat shock proteins also reported to be required for the resistance mediated by R proteins^8^. These findings on CMD resistance in cassava (i) further highlight the complexity of cassava mosaic disease and the fact that current qualitative methods of CMD phenotyping do not provide enough information to decipher the molecular relationship between genotype and phenotype, given that *CMD2* gene was not differentially expressed. These findings are similar to what has been previously reported for CBSD^58^. (ii) In addition, this could also be attributed to the fact that the *CMD2* gene is not expressed on any of the five cassava tissues in this study (as seen in **Supplementary Figure 9b - right side**), as previously speculated by Adenike et al^59^. Given that previous studies on CMD resistance highlighted a single source of monogenic resistance in the cassava genepool^60^, especially in Africa. The findings in this study provide the foundation for a more in-depth, quantitative understanding of resistance and support the long term goal of diversifying the sources of resistance, given the precarious nature of single gene resistance^9,10^.

## Conclusion

While transcriptomics has relied on contrasting individuals, our study provides a population-based resource, unique to previously described available transcriptomics cassava resources. Resources including genomics, transcriptomics, metabolomics, epigenomics and proteomics to support cassava improvement and intensification have been previously described^15,32,61–71^. In this study, we characterized the transcriptomics of 150 accessions across five different tissue types. The expression dataset highlights the similarity in expression pattern between tissue types, indicating that tissue types that are physically closely related seem to have similar expression patterns. We detected 19,445 DEGs with 8,820 DEGs unique across tissue types comparisons and further characterisation detected 18 modules, in which the HCN cassava root regulation transporter, the galactose transporter and the plant defence gene against cassava mosaic virus and its insect vector whitefly, were all located in the yellow module, based on previous characterization in cassava. The yellow module is widely involved in developmental processes and highlights important regulatory genes in cassava. These 307 genes in the yellow module represent important genes for studying physiology and developmental characters for cassava improvement. Given that they were highly enriched in biosynthesis of secondary metabolites and metabolic pathway processes. We provided further insight on CMD resistance and highlighted sets of 27 genes that were involved in the plant-pathogen interaction pathway. Our study suggests a potential path to sources of CMD resistance diversification and provides a queryable cassava expression atlas that will serve as a valuable and novel population-based resource to the cassava research community. These resources can be used to develop a biological information driven genomic selection (GS) framework to further improve prediction accuracies, especially for disease traits, which could leverage spatial and temporal control of plant pathogen response to facilitate breeding for crop improvement in cassava. Other applications include differential gene expression analysis to quantitatively evaluate transcript levels between tissue types or contrasting individuals of interest and performing expression quantitative trait loci to identify genomic loci explaining variation in expression levels of mRNAs for traits of interest. In the future, a validation study of the Manes.10G147700 gene using analysis of promoters or knock-outs would be required to understand its relationship with the CMD phenotype. The combination of these approaches would provide information on the specific functioning of Manes.10G147700 to CMD inoculation at different developmental stages.

## Methods

### Tissue sample collection, RNA extraction, RNA-seq library preparation and Illumina sequencing

Tissue samples were collected from 150 accessions across five tissues at a cassava experimental field at National Crops Resources Research Institute (NaCRRI) Kampala, Uganda. These accessions are part of the genetic populations of NaCRRI^10^. For most of the accessions, samples were collected from leaf, stem, fibrous root, storage root and flower tissues; while for accessions that were not flowering, apical meristem tissue was collected in place of flowers. RNA was extracted using TRIzol by Invitrogen protocol, in which hot borate and lithium chloride were used to extract RNA from tissue samples of leaf, flower, stem, fibrous and storage root^72^. 3’RNA-seq libraries were prepared according to the method described by Kremling et al^18^. Briefly, libraries were prepared robotically, using 96-well plates from 500 ng total RNA on an NXp liquid handler using QuantSeq FWD kits. Post-PCR cleanup was performed and libraries were pooled to 96-plex according to the QuantSeq protocol. Molar concentrations were calculated for each pool and sequenced using 90 nucleotide single-end read Illumina TruSeq primers on an Illumina NextSeq 500 with v2 chemistry in the year 2015 at the Cornell University Sequencing facility. Normalized gene-leve counts and raw sequencing datasets were deposited on cassavabase.org, hosted at ftp://ftp.cassavabase.org/Cassava3primeRNAseqRawReads/. Raw sequencing data were submitted to NCBI sequence reads archive (SRA) with the following details (BioProject: PRJNA737128) and can be accessed via http://www.ncbi.nlm.nih.gov/bioproject/737128.

### 3’RNA-seq dataset processing

Fastq files for each sample were processed using Trimmomatic^73^ (version 0.32) to remove the first 12 bp and remnants of the Illumina Truseq adapter from each read according to kit maker recommendation. The STAR aligner^74^ (version 2.7) was used to align reads against cassava version 6.1 (https://phytozome.jgi.doe.gov/pz/portal.html#!info?alias=Org_Mesculenta) genome annotation using parameters that included (-outFilterMultimapNmax 10; -outFilterMismatchNoverLmax 0.04; - outFilterIntronMotifs RemoveNoncanonicalUnannotated), allowing reads to map in 10 locations, with at most 4% mismatches, and filtering out all non-canonical intron motifs. HTSeq^75^ (version 0.11.2) default settings were used to obtain gene-level counts, which were then normalized using the counts per million mapped reads (CPM) method implemented in the EdgeR^23^ (version 3.26.8) package in R version 3.6.3 (2020-02-29). The normalized CPM expression digital values were used for visualization in a cassava expression atlas^76^ (https://cea.sgn.cornell.edu/expression_viewer/input) hosted on https://cassavabase.org (https://cassavabase.org) (see Supplementary Fig. S11 for data processing schema). Obtained gene-level counts from HTSeq^75^ were normalized using variance stabilizing transformation^77^ (vst) method in DESeq2^77^ (version 1.24.0) for downstream analysis including principal component analysis (PCA) and weighted gene co-expression network analysis (WGCNA)^78^.

### Cassava Expression Atlas data availability

The cassava expression atlas graphical interface was implemented on https://cassavabase.org to allow for the interactive visualization and exploration of tissue-specific patterns and discovery of trends in a population-based transcriptomics dataset^76^. The implementation uses CPM normalized read counts obtained from HTSeq^75^ (version 0.11.2), functional gene annotations from Phytozome (version 10.3), and analysis of correlation of genes using the *cor* function in R 3.4.2 (R. Core Team, 2015). The expression atlas has four main features: expression cube, expression images, heatmap and scatter plot. The expression cube enables gene discovery based on gene expression patterns across accessions and tissue types, with the ability to display genes with expression correlated to that of your gene of interest across the dataset. The expression images are whole cassava plant images showing tissue-specific expression patterns of selected or newly discovered genes across selected accessions. The heatmap is created to visualize tissue-specific expression patterns for selected genes across selected accessions. The scatter plot feature visualizes expression of any two samples. The cassava expression atlas can be found at: https://cea.cassavabase.org and additional details can be found in the https://cassavabase.org manual

### Differential gene expression and weighted Gene Co-expression Network analysis

Differential expression and statistical analysis were carried out using the DESeq2^77^ (version 1.24.0) R package. DESeq2 uses raw read counts as input, corrects for library size and accounts for sequencing depth using vst normalization. To account for multiple testing corrections, p-values were adjusted using Benjamini-Hochberg^79^ testing procedure and a false discovery rate adjusted p-value of <0.05 was used as a threshold for significantly differentially expressed genes (DEGs) for each differential expression analysis done in this study. As a quality assessment step, we profiled the expression of Rubisco as a positive control similar to housekeeping genes used in qPCR^80^. To assemble DEGs for downstream analysis, first we retrieved the detected DEGs for different pairwise tissue comparisons based on adjusted p-value of <0.05 and log2 fold change (log2FC) of >|1|. We then combined (union) all detected DEGs (19,445) from all comparisons and kept a total of 8,820 unique DEGs for downstream analysis. Previously reported resistance genes with cassava genome annotation versions earlier than version 6 were identified by BLAST search on phytozome (https://phytozome.jgi.doe.gov/pz/portal.html#!search?show=BLAST) using cassava genome version 6 annotation. Variance stabilizing transformation normalized gene-level counts from HTSeq for the 8,820 selected unique DEGs were used to infer co-expression gene network modules using the WGCNA^78^ R package with power-law coefficient **β** selected using the soft-thresholding method and a hierarchical tree cut algorithm used in detecting the co-expression modules.

### Gene Ontology (GO) analysis

Genes identified in different modules from WGCNA were characterized for GO terms including biological processes, cellular components, and molecular functions using over-representation Fisher’s exact test with Bonferroni multiple testing correction p-value of < 0.05 in PANTHER version 16.0 (Released 2020/07/28) as previously described^10^. The gene enrichment analysis was performed on the *Arabidopsis thaliana* GO database (GO Ontology database DOI: 10.5281/zenodo.4081749 Released 2020-10-09)^31^ using corresponding cassava genome version 6.1 annotation IDs^32^ as previously described^10^. DEGs identified from differential expression analysis of CMD susceptible and resistant accessions were subjected to KEGG pathways enrichment analysis ^29,30^. KEGG was used because it was easy to overlay identified genes on pathway maps available on the platform (*Database last updated: January 18, 2021*).

### Partitioning proportion of genetic variance explained by markers on the DEG selected regions

Using parametric multiple kernel mixed model^81^ as previously described for cyanide^33^ and other traits in cassava, we calculated the heritability contribution of the SNP markers found within the regions (with 53335 bp maximum transcript length) of the selected 8,820 DEGs for traits fresh root yield (FYLD), vigor, root number (RTNO), dry matter (DM), mean CMD (MCMD), and mean CGM (MCGM). The variance explained by the markers in these regions was compared with a random set of markers of similar size and distribution across the genome. To ensure precise estimation of the proportion of variance explained, LD was controlled by removing markers that are in high LD (r2 >= 0.9) with the 11,399 SNPs in the DEG regions. We estimated the variance components using the ‘emmremlMultikernel’ function implemented in the R package EMMREML^82^. The multikernel model is represented in matrix notation: 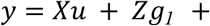 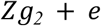, where *y*is the vector of the best linear unbiased prediction (BLUP) for each trait, and *X* is a vector of ones, representing the intercept. *u*is the genetic mean effect of DEG SNPs on a trait, and *Z* is the design matrix linking observations to individuals. *g*_*1*_ *and g*_*2*_are the genetic variance components for SNPs in the DEG regions and sampled SNPs of equal size and distributions across the genome, respectively. Where *g*_*1*_ ~ *N*(*0*, *GRM*_*D*_*σ*^*2*^) and *g*_*2*_ ~ *N*(*0*, *GRM*_*S*_*σ*^*2*^) have known variance structure, calculated using DEG SNPs (*GRM*_*D*_) or sampled SNPs (*GRM*_*S*_) and *e* is the residuals variance. The phenotype and genotype dataset used were sourced from Okeke et al. and included 750 accessions and 76,453 SNP markers after a minor allele frequency filtering threshold of 0.01, below which SNPs were removed^3^. These historical datasets are from multiple trials conducted at the International Institute of Tropical Agriculture, Ibadan, Nigeria, as a part of their genetic gain population and represented clones selected between 1970s through 2007^83^.

## Acknowledgements

The authors appreciate Susan McCouch for providing helpful discussion and feedback. We acknowledged that the dataset used in this study was generated, sequenced and deposited on https://cassavabase.org ftp site (ftp://ftp.cassavabase.org/Cassava3primeRNAseqRawReads/) by the National Crops Resources Research Institute, Ugandan. This work was partially supported by the NEXTGEN Cassava project, through a grant to Cornell University by the Bill & Melinda Gates Foundation (Grant INV-007637 http://www.gatesfoundation.org) and the UK’s Foreign, Commonwealth & Development Office (FCDO).

## Author contributions statement

Designed the research: ACO, EW, RP, GB, JLJ, LM. Performed the research: ACO. Contributed to Cassava Expression Atlas: ACO, GB, NM, AP. Contributed data: LN, EW, RK. Wrote the paper: ACO. Supervised the research: LM. Contributed to the manuscript editing: RP, GB, LM, EW, NM, AP, LN. All authors reviewed the manuscript and gave final approval for the publication.

## Additional Information

Correspondence and requests for materials should be addressed to ACO or LAM.

## Supplementary Information

Supplementary Figure S1 - S11

Supplementary Table S1: Variance-stabilized transformed reads count.

Supplementary Table S2: Counts per million mapped reads (CPM) normalized reads count.

Supplementary Table S3: Raw reads count.

Supplementary Table S4: Differentially expressed genes (DEGs) across all contested tissue types.

Supplementary Table S5: Unique 8,820 differentially expressed genes (DEGs).

Supplementary Table S6: CMD differentially expressed genes (DEGs) for Mkumba versus NASE14.

Supplementary Table S7: CMD DEGs observed in Plant-pathogen interactions pathway.

Supplementary Table S8: List of differentially expressed genes WGCNA modules.

Supplementary Table S9: Identified PANTHER gene ontology terms.

Supplementary Table S10: Detected modules GO terms enrichment descriptions.

Supplementary Table S11: Resistant genes previously reported by Allie et al., 2014.

