## Supplementary Figures 1 - 11 for "A population based expression atlas provides insights into disease resistance and other physiological traits in cassava (*Manihot esculenta* Crantz)"

**Supplementary Figure 1.** Cassava expression atlas (CEA).

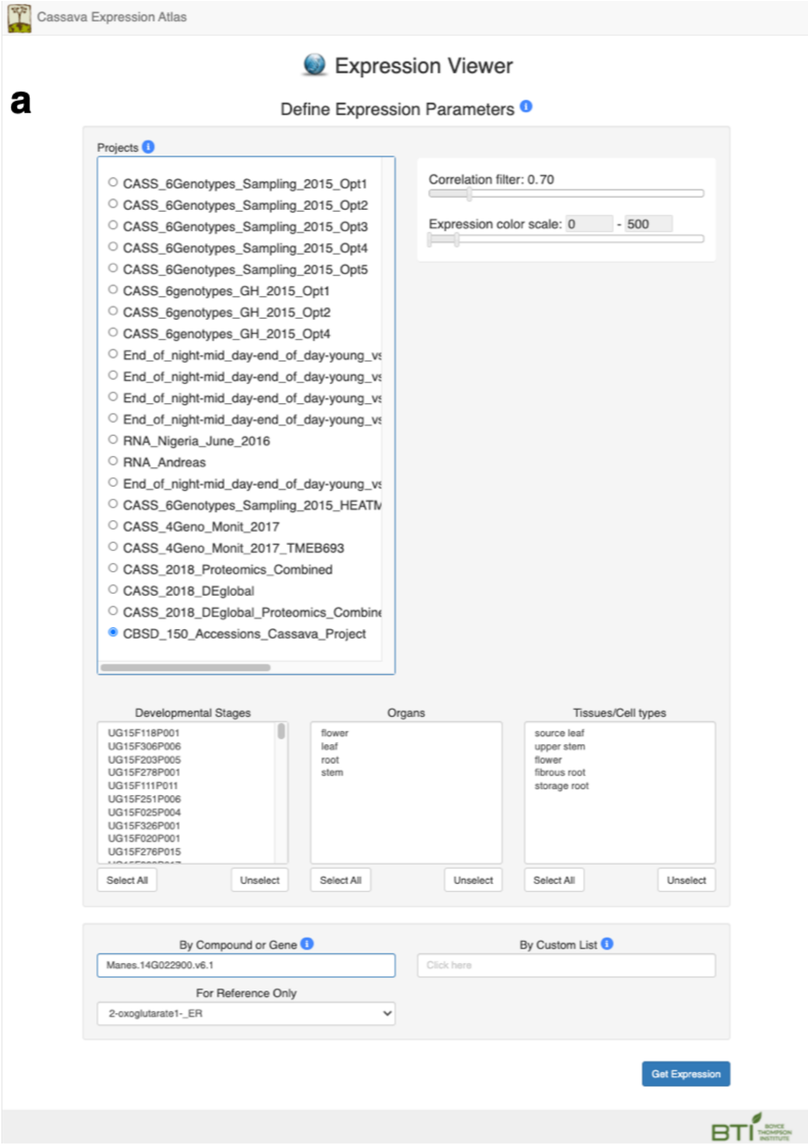

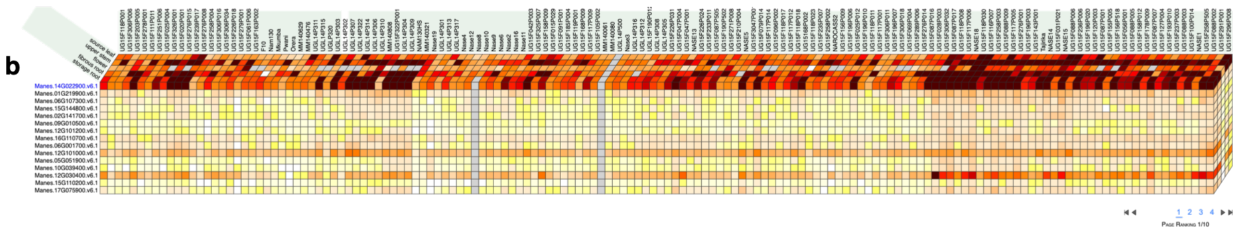

To visualize the CEA for any gene and specific accession or sets of accessions, select project name (CBSD_150_Accessions_Cassava_Project) from the home page (https://cea.sgn.cornell.edu/expression_viewer/input), select accessions, organs and tissue types from their “textboxes” below the project name. Then enter the gene name in the “By Compound or Gene" textbox and click the “Get Expression” button. (A) Home page of CEA showing project, accessions, organs, and tissues selection panel. (B) CEA cube showing expression of 150 accessions across 5 tissues for Manes.14G022900 gene and other genes correlated with the gene of interest with correlation coefficient of 0.65 and above.

**Supplementary Figure 2.** Differential expression analysis.

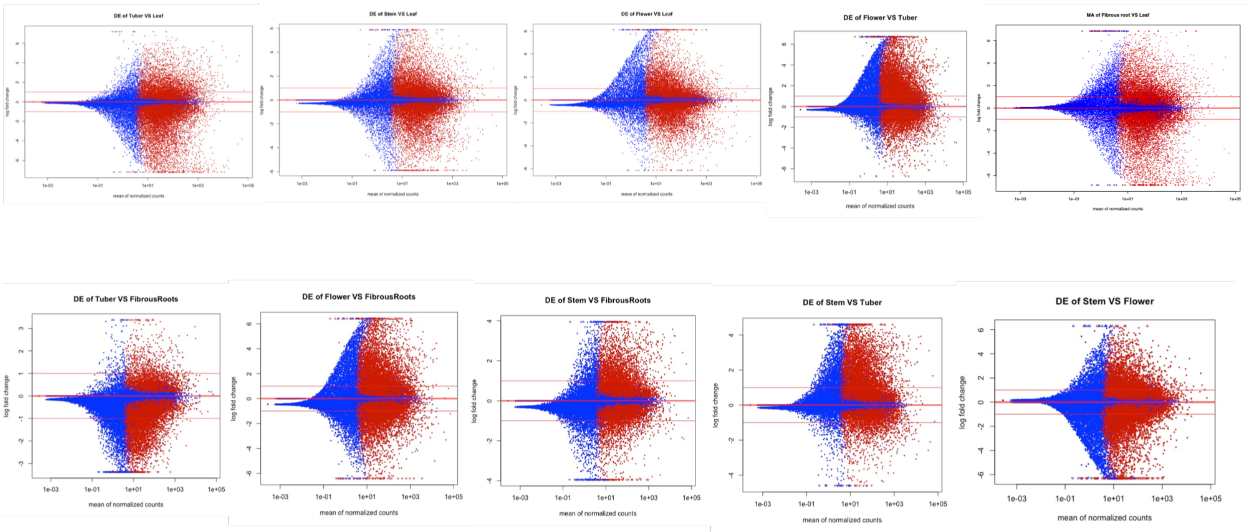


Differential gene expression analysis plots showing differentially expressed genes between contrasted tissue types. Here “Tuber” is referred to as “Storage root”. Red dots represent genes that were differentially expressed based on adjusted p-value of < 0.05. The blue dots represent genes that are not differentially expressed. Log2 fold change was plotted on the y-axis, while mean of normalized counts was plotted on the x-axis

**Supplementary Figure 3.** Volcano plots showing the differentially expressed gene across all contrasted tissue types.

a

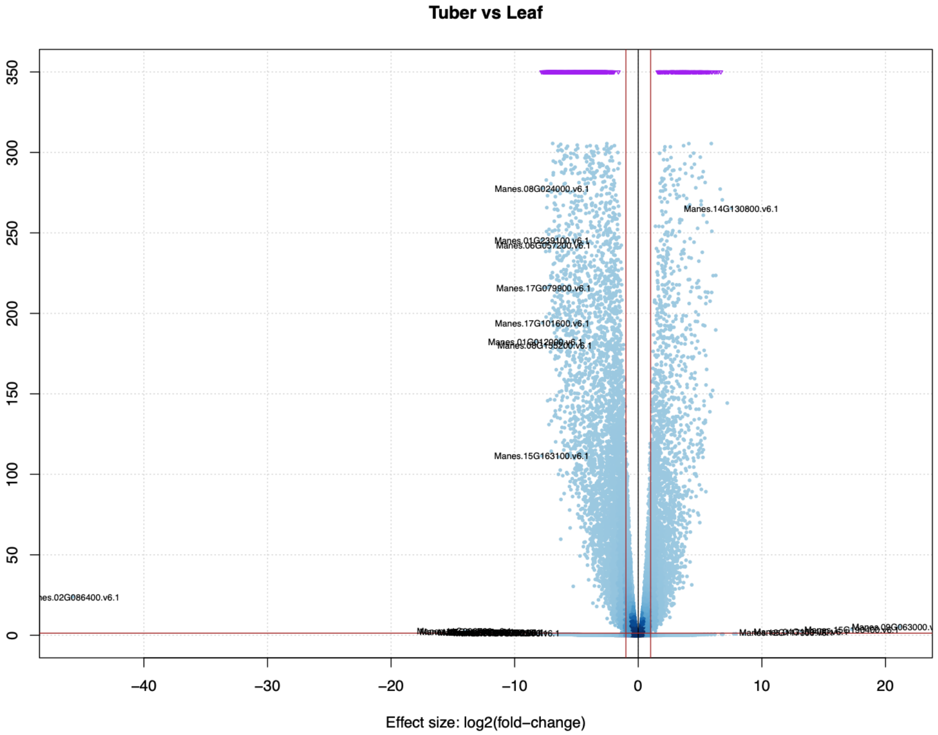


b

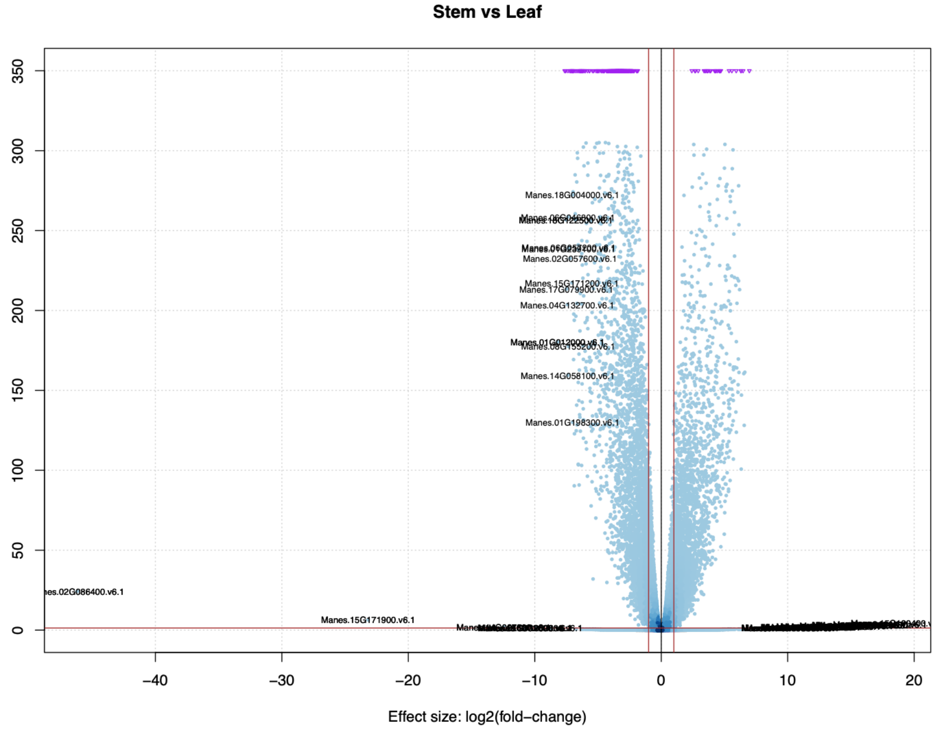

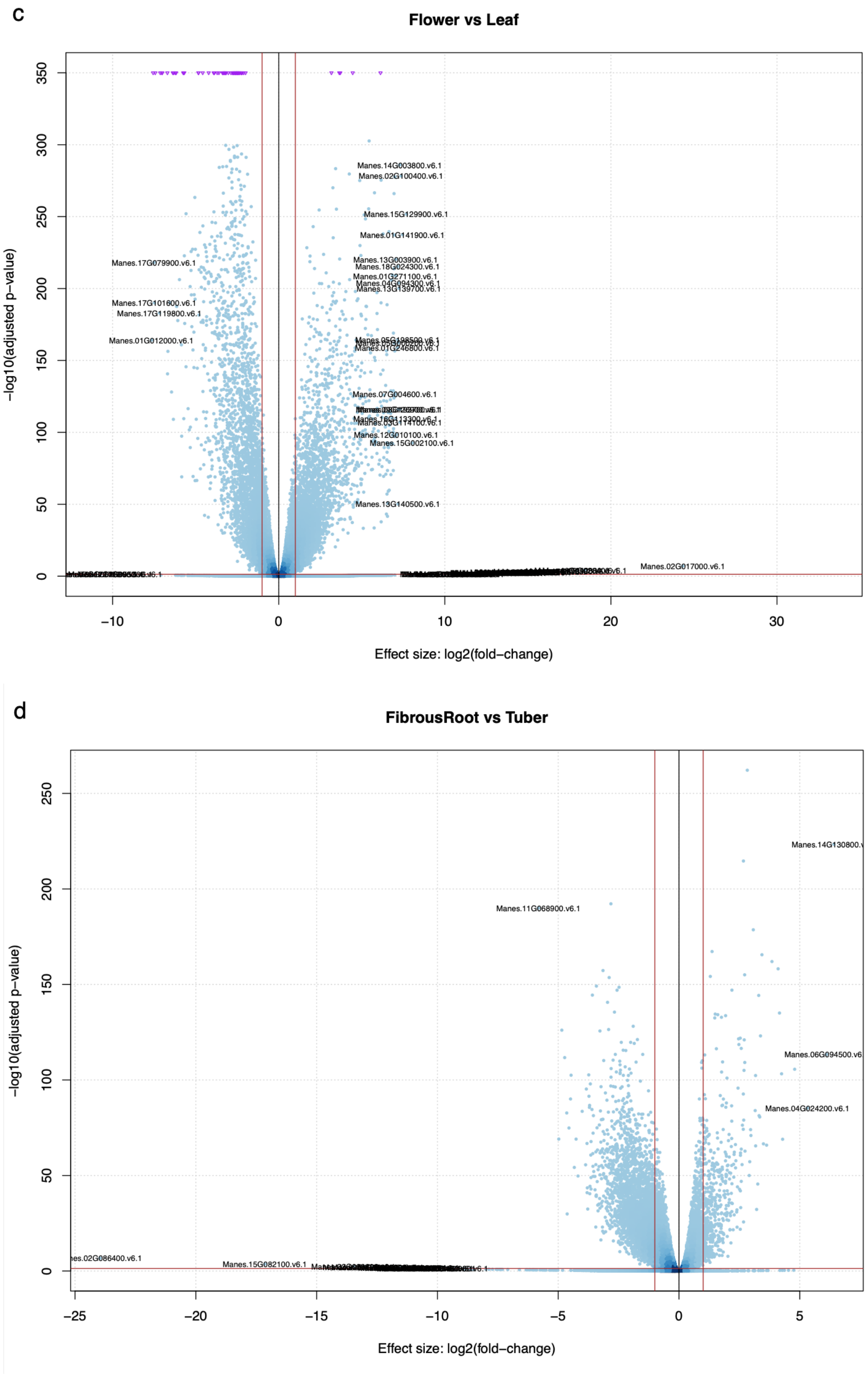
c
Flower vs Leaf
−log10(adjusted p−value)
Effect size: log2(fold−change)
Manes.14G003800.v6.1
Manes.02G100400.v6.1
Manes.15G129900.v6.1
Manes.01G141900.v6.1
Manes.17G079900.v6.1
Manes.13G003900.v6.1
Manes.18G024300.v6.1
Manes.01G271100.v6.1
Manes.04G094300.v6.1
Manes.13G139700.v6.1
Manes.17G101600.v6.1
Manes.17G119800.v6.1
Manes.01G012000.v6.1
Manes.01G246800.v6.1
Manes.07G004600.v6.1
Manes.03G114100.v6.1
Manes.12G010100.v6.1
Manes.15G002100.v6.1
Manes.13G140500.v6.1
Manes.02G017000.v6.1
d
FibrousRoot vs Tuber
−log10(adjusted p−value)
Effect size: log2(fold−change)
Manes.11G068900.v6.1
Manes.04G024200.v6.1
Manes.15G082100.v6.1

e

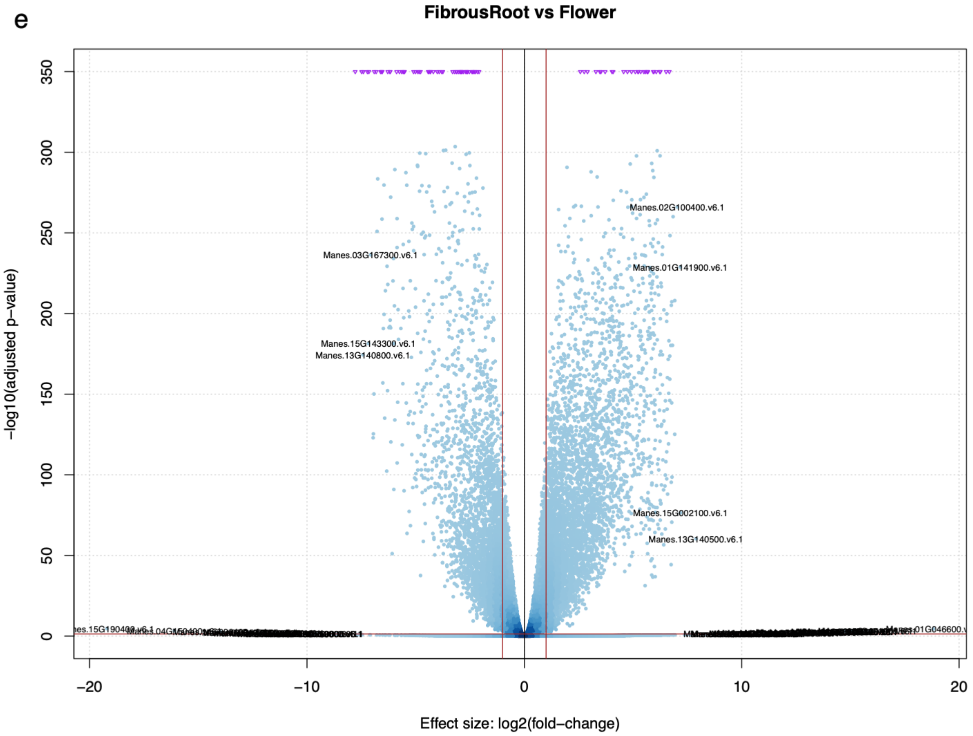

FibrousRoot vs Flower
−log10(adjusted p−value)
Effect size: log2(fold−change)
Manes.02G100400.v6.1
Manes.03G167300.v6.1
Manes.01G141900.v6.1
Manes.15G143300.v6.1
Manes.13G140800.v6.1
Manes.15G002100.v6.1
Manes.13G140500.v6.1


f

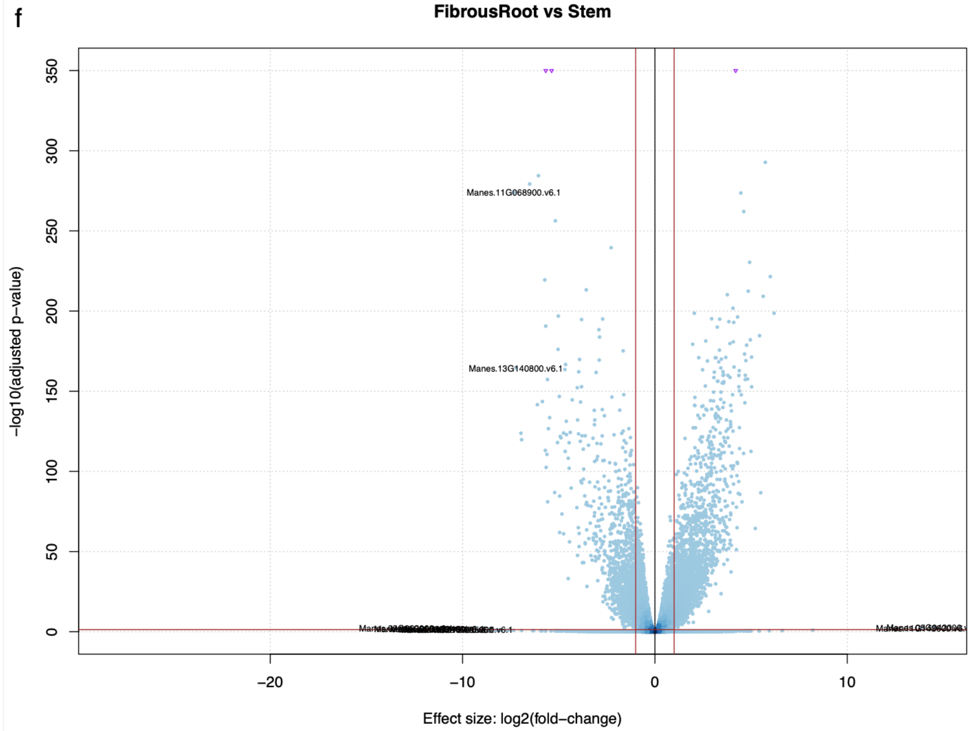

FibrousRoot vs Stem
−log10(adjusted p−value)
Effect size: log2(fold−change)
Manes.11G068900.v6.1
Manes.13G140800.v6.1

g

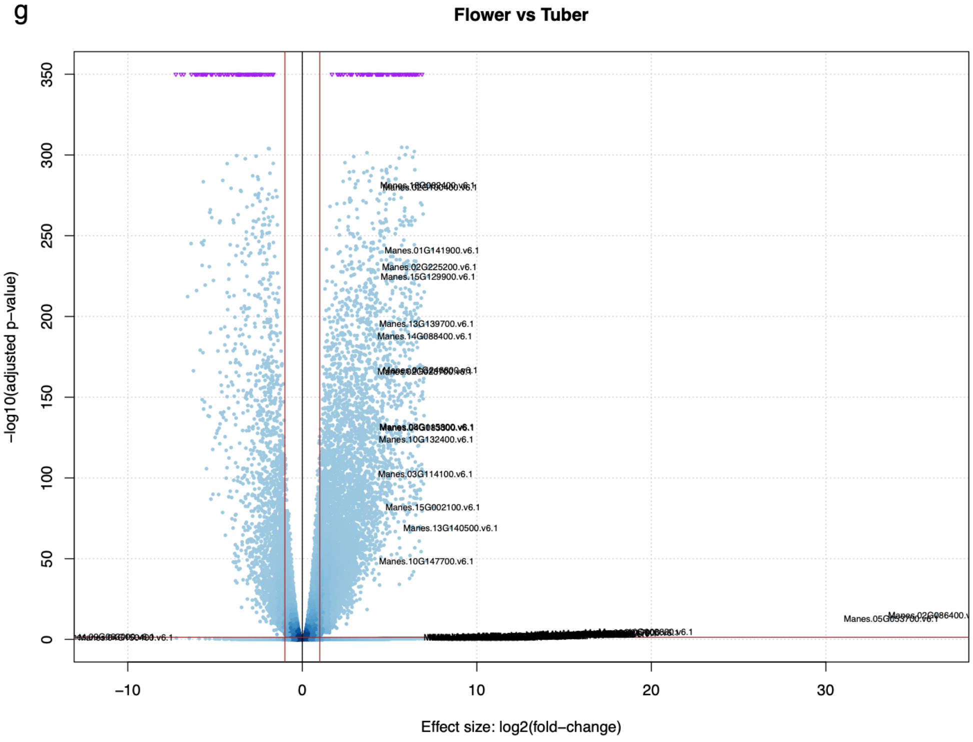
Flower vs Tuber
-log10(adjusted p-value)
Effect size: log2(fold-change)
Manes.01G141900.v6.1
Manes.02G225200.v6.1
Manes.15G129900.v6.1
Manes.13G139700.v6.1
Manes.14G088400.v6.1
Manes.10G132400.v6.1
Manes.03G114100.v6.1
Manes.15G002100.v6.1
Manes.13G140500.v6.1
Manes.10G147700.v6.1

h

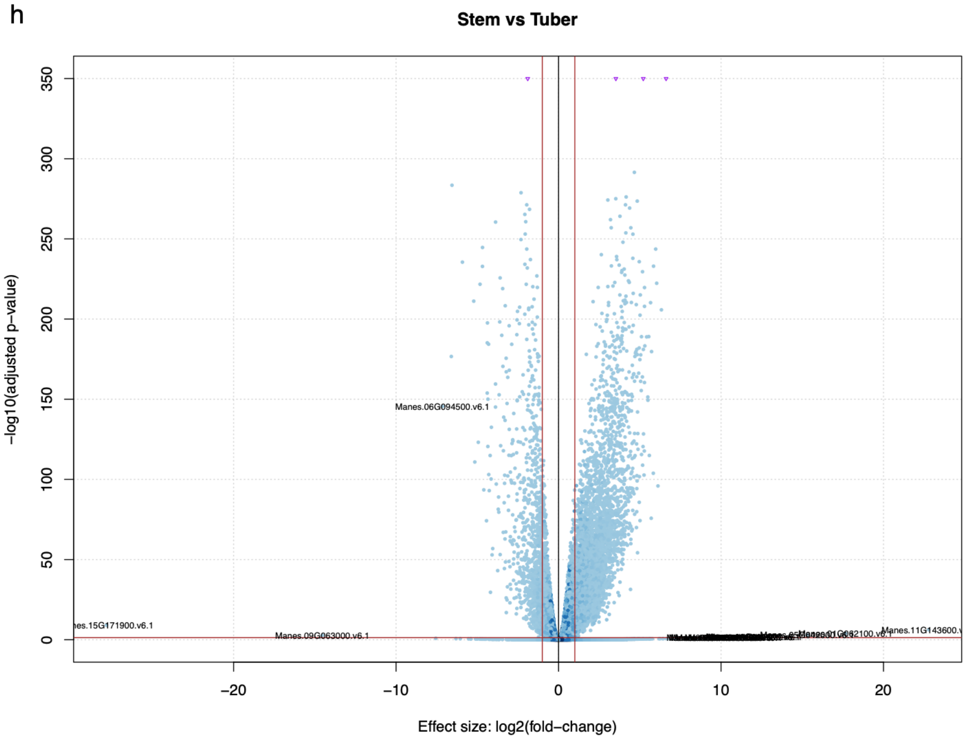
Stem vs Tuber
-log10(adjusted p-value)
Effect size: log2(fold-change)
Manes.06G094500.v6.1
Manes.09G063000.v6.1

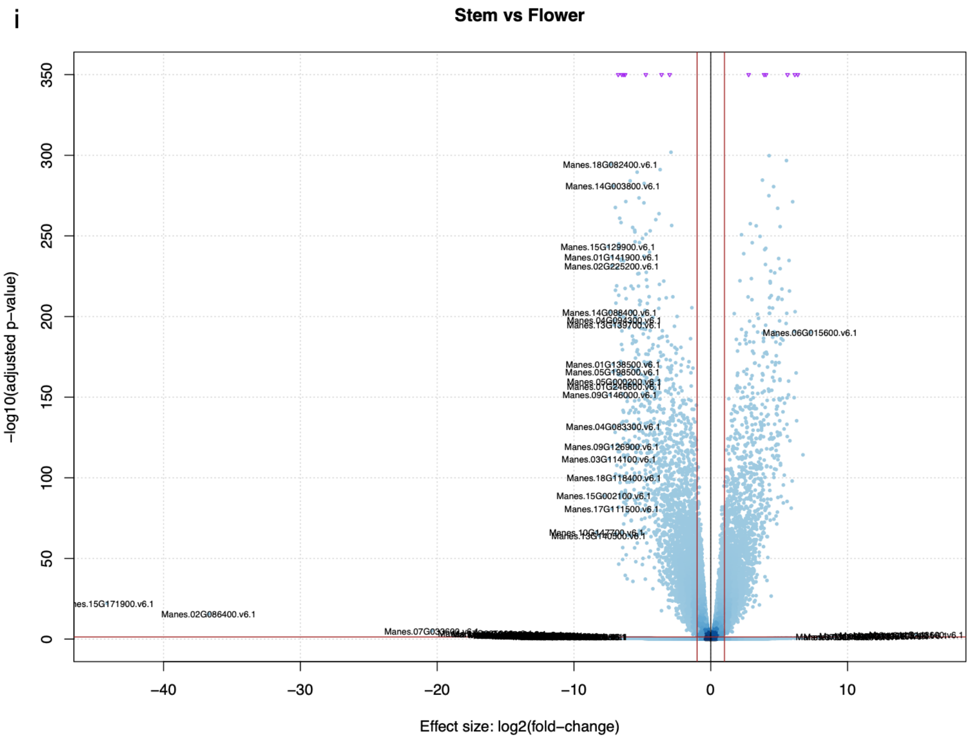


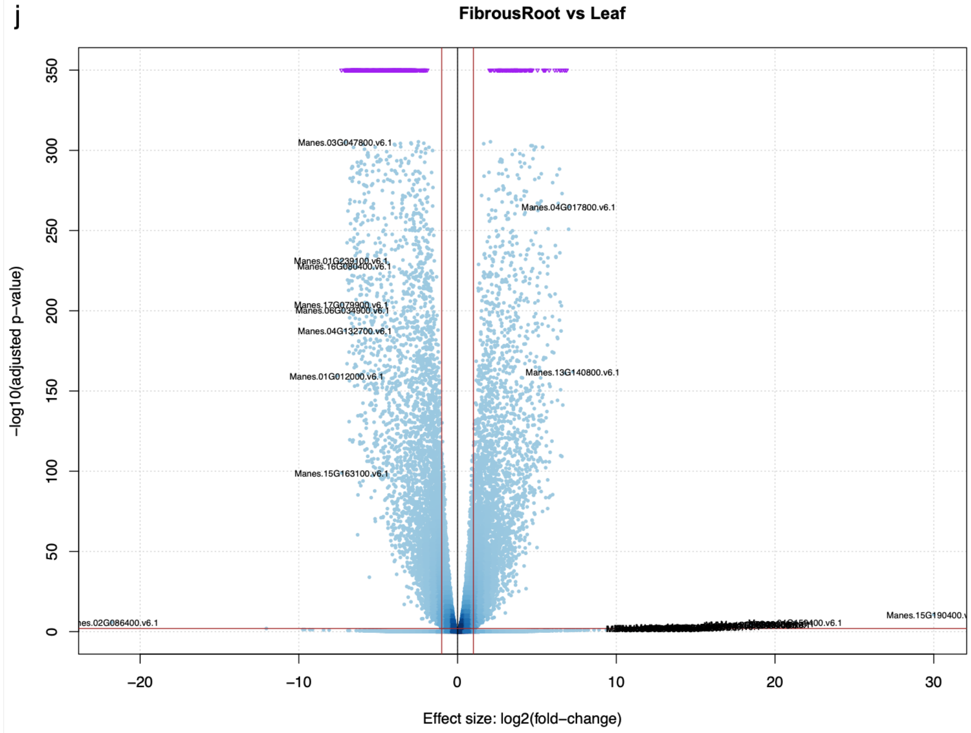


Volcano plots of differentially expressed genes (DEGs) showing up- and down-regulated DEGs. Negative log10 of the adjusted p-value was plotted on the y-axis, while effect size (log2 fold change) was plotted on the x-axis. The vertical red lines were log2 fold change threshold of > |1|. The horizontal line is the adjusted p-value threshold < 0.05. (A) storageRoot vs leaf. (B) stem vs leaf. (C) flower vs leaf. (D) fibrousRoot vs storageRoot. (E) fibrousRoot vs flower. (F) fibrousRoot vs stem. (G) flower vs storageRoot. (H) Stem vs storageRoot. (I) stem vs flower. (J) fibrousRoot vs leaf. Gene names printed on the plots are those with the highest effect size based on "log2 fold change" across contracted tissue. Here "Tuber" is referred to as "storage root".

**Supplementary Figure 4.** Volcano plot and expression atlas display for Rubisco Activase protein and Rubisco methyltransferase family protein.

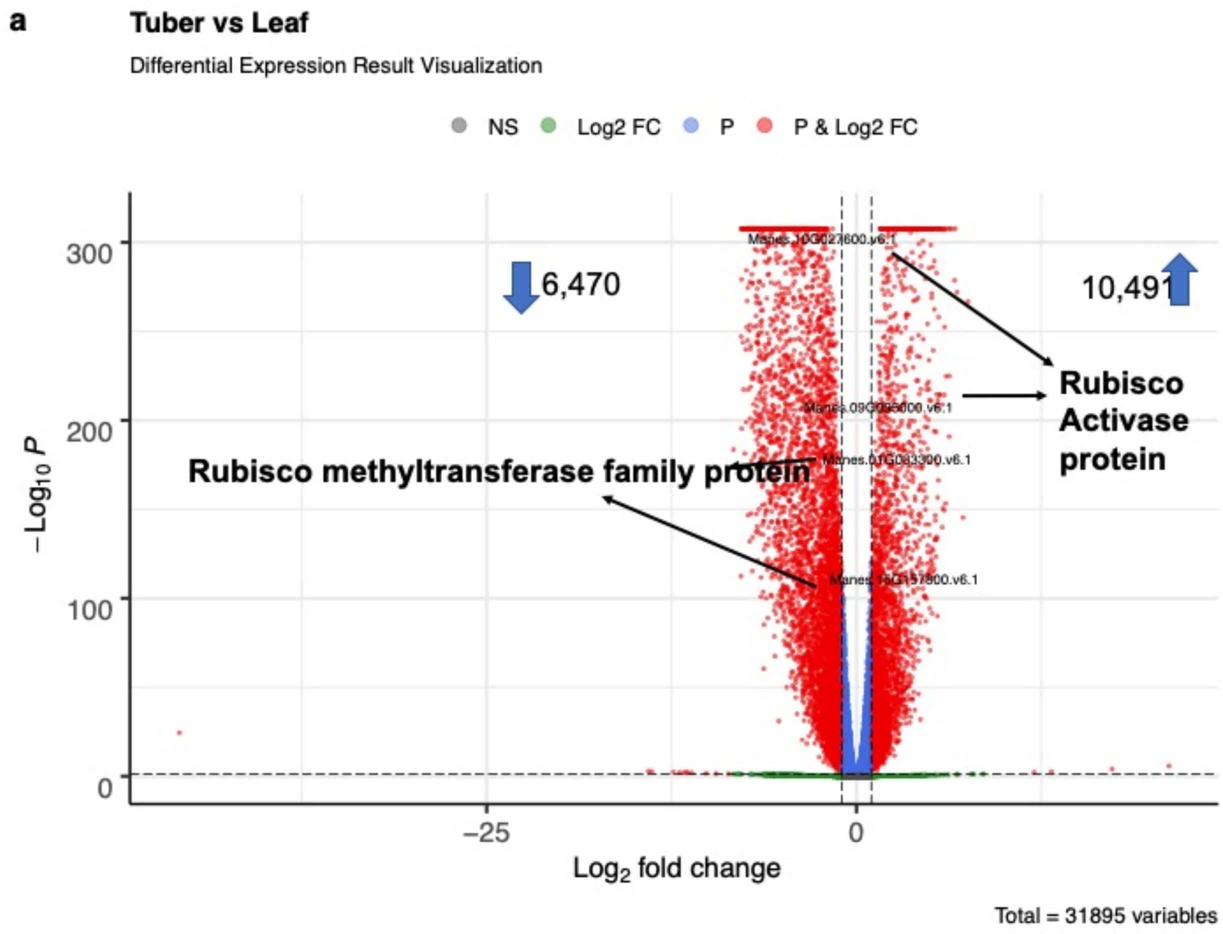

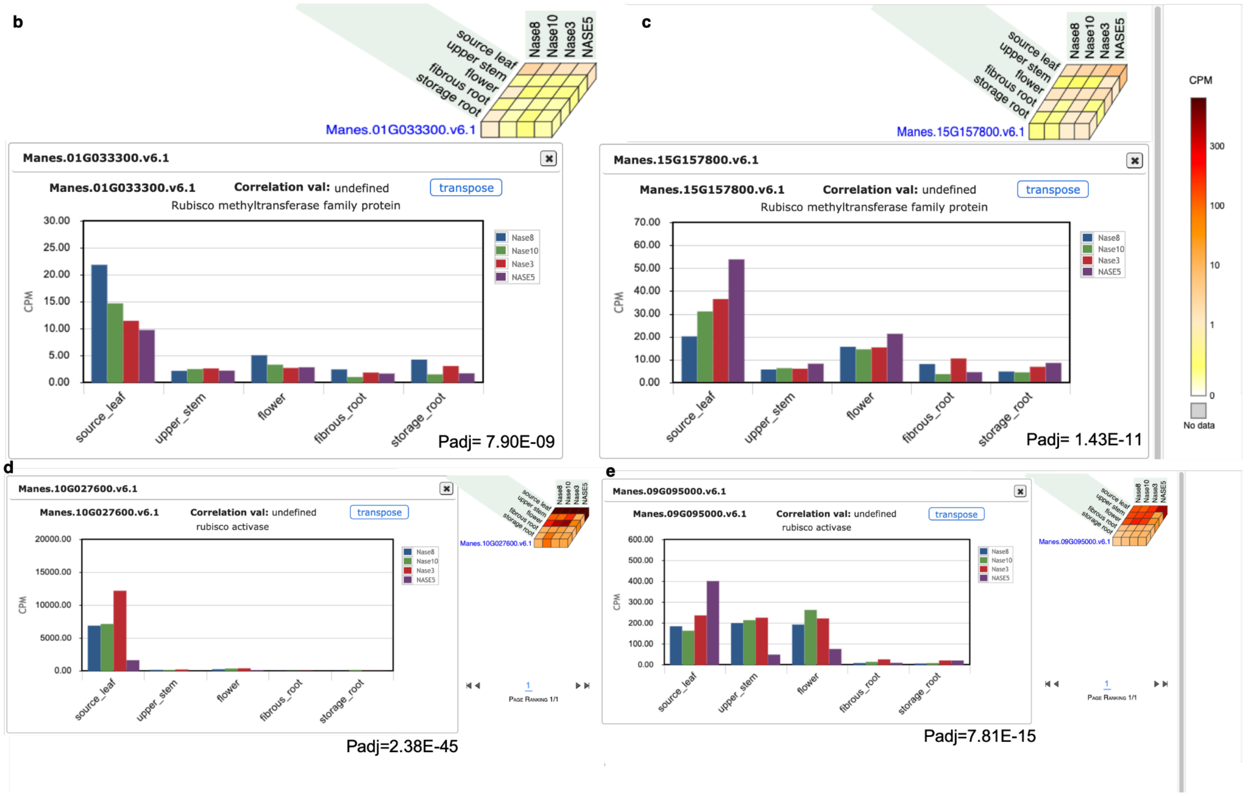


(A) Volcano plot showing the differentially expressed gene between contrasted tissue of storage root vs leaf. Here "Tuber" is referred to as "storage root". Gene names printed on the plots were selected for their visualization on cassava expression atlas (CEA). Negative log10 of the adjusted p-value was plotted on the y-axis, while effect size (log2 fold change) was plotted on the x-axis. The vertical red lines were log2 fold change threshold of > |1|. The horizontal line is the adjusted p-value threshold < 0.05. (B-C) Expression atlas cube [upper panel] and barplot [lower panel] showing the expression of five tissue types on four accessions for Manes.01G033300 and Manes.15G157800 gene, a Rubisco methyltransferase family protein. The gene was observed to be significantly differentially expressed between cassava leaf and storage root as expected. (D-E) Expression atlas cube [upper panel] and barplot [lower panel] showing the expression of five tissue types on four accessions for Manes.10G027600 and Manes.09G095000 gene, a Rubisco activase protein. Other genes correlated (r2 >= 0.65) to the gene of interest were listed below in the expression cube. These genes were observed to be significantly differentially expressed between cassava leaf and storage root as expected. Rubisco methyltransferase family proteins are enzymes involved in the major step of carbon fixation, a process that converts carbon dioxide from the atmosphere into energy rich molecules such as glucose; Rubisco activase proteins are chloroplast proteins required for light activation of rubisco[1] and jasmonate-induced leaf senescence[2].

**Supplementary Figure 5.** Differential expression analysis between Mkumba [CMD resistant ] vs Nase14 [CMD susceptible].

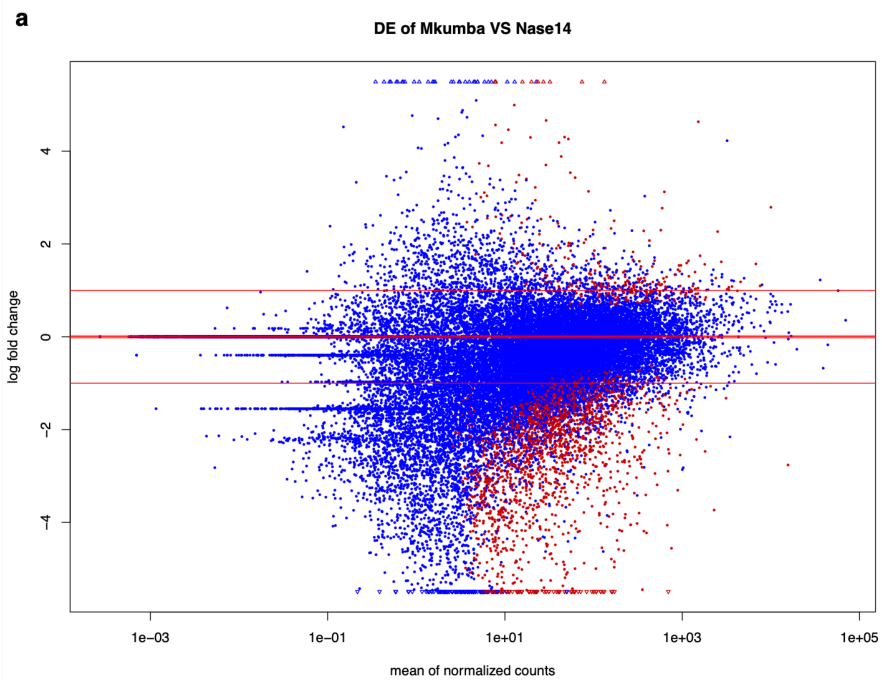


Plot showing differentially expressed genes based on adjusted p-value of > 0.05. The blue dots represent genes that are not differentially expressed. Log2 fold change was plotted on the y-axis, while mean of normalized counts was plotted on the x-axis.

**Supplementary Figure 6.** Scale free topology (SFT) criterion for identified DEGs weighted gene co-expression network analysis.

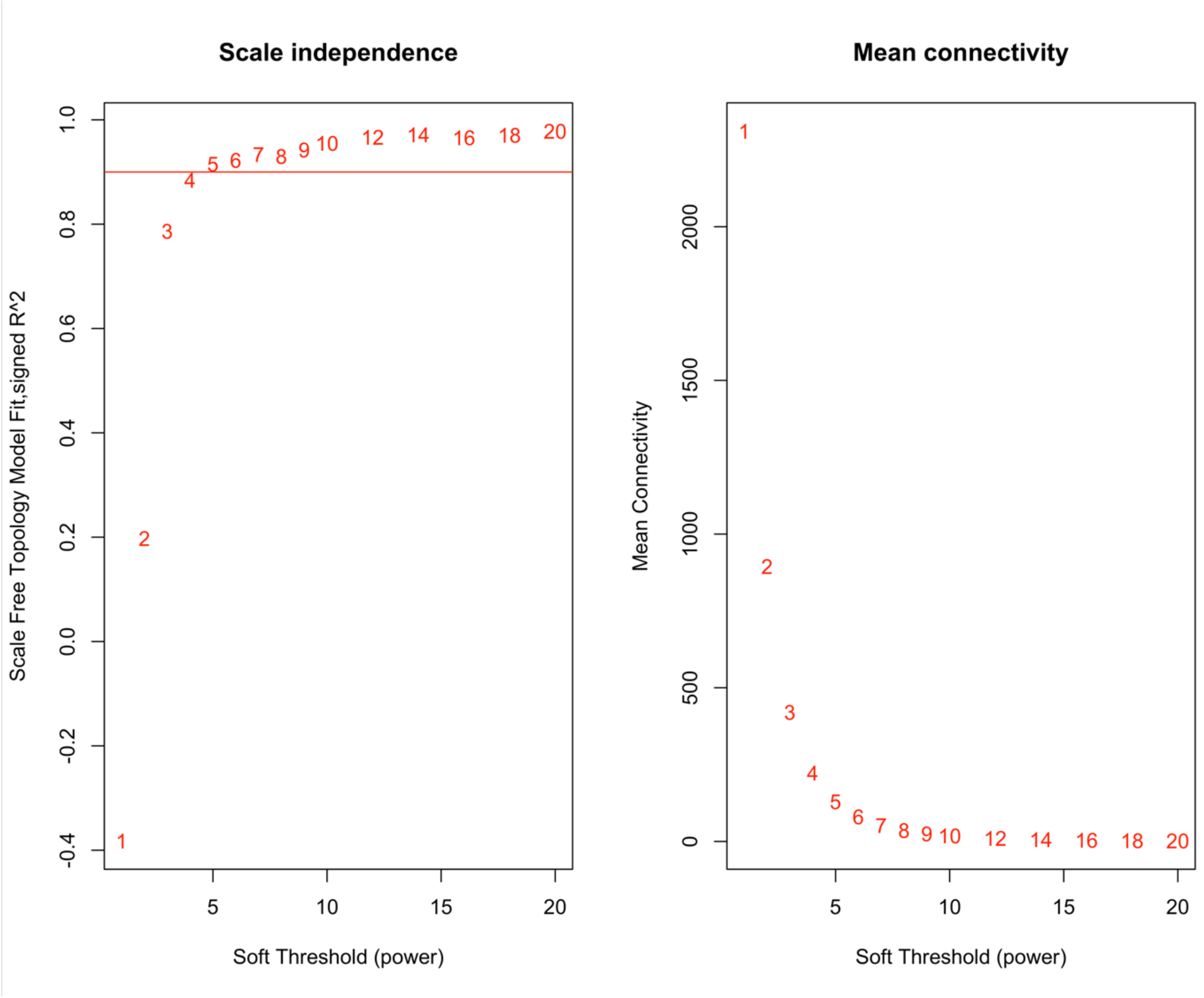


SFT plot for choosing the power beta for the unsigned weighted correlation network. Left plot: the SFT index R2 (y-axis) as a function of different powers of beta (x-axis). The value of beta chosen was 7 and has a corresponding R2 value of 0.93. Right plot: the mean connectivity on the y-axis is a strictly decreasing function of the power beta, on the x-axis.

**Supplementary Figure 7.** Dendrogram with module colors for detected 8,820 DEGs and PANTHER GO-Slim Molecular Function categorization for genes in the yellow module.

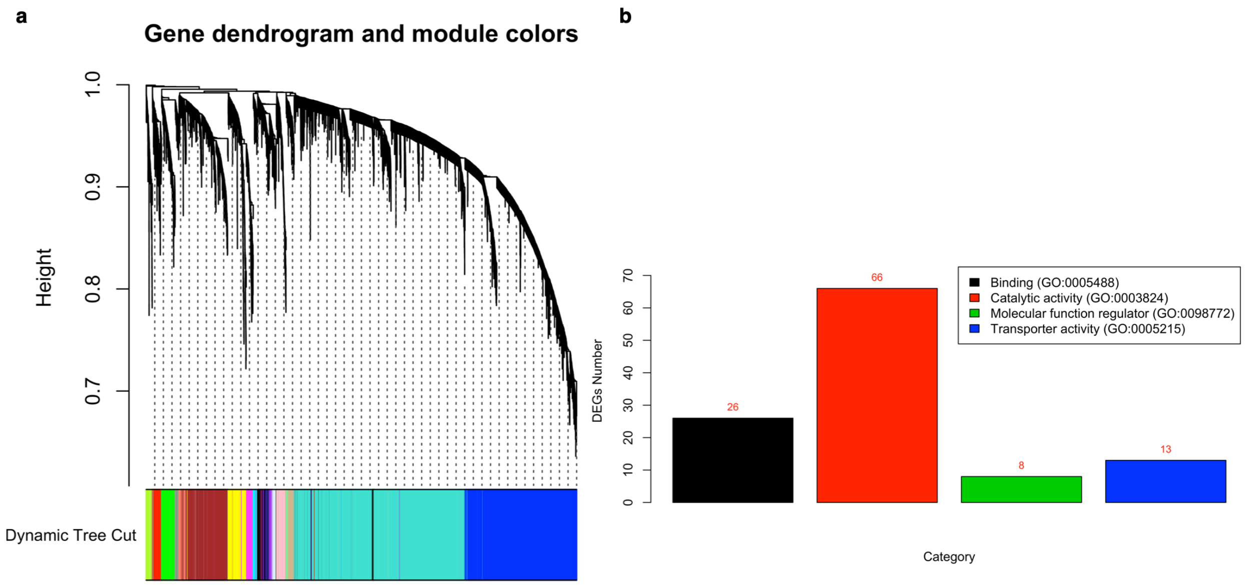


(A) Dendrogram with module colors. (B) Barplot showing PANTHER GO-Slim Molecular Function categories of genes in the yellow module. One hundred and thirteen (113) genes were characterized based on GO-Slim Molecular Function while 194 are uncharacterized in the molecular function category.

**Supplementary Figure 8.** PANTHER Gene Ontology categorization of genes in the yellow module.

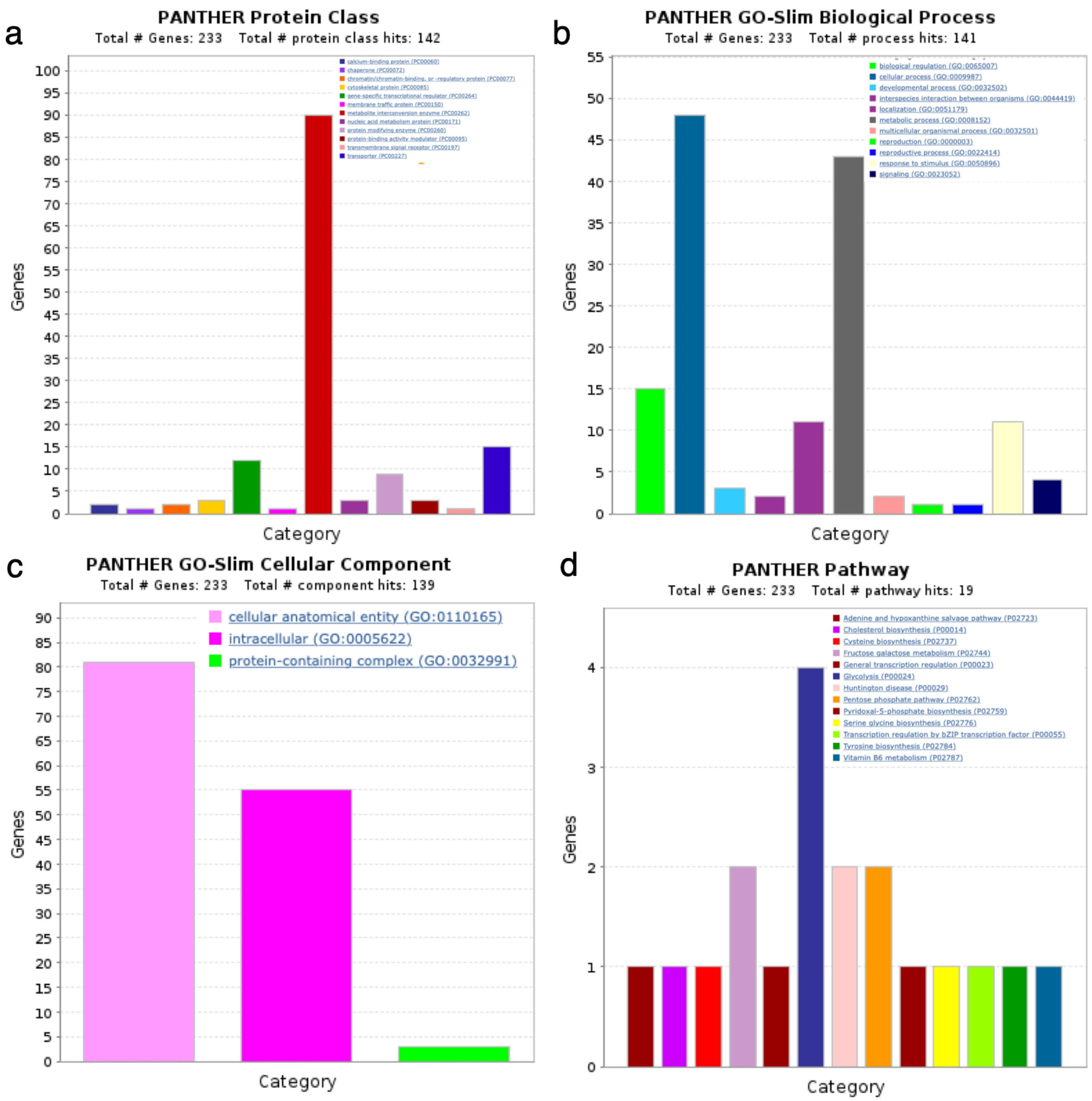


(A) Barplot showing PANTHER Protein class categories of genes in the yellow module. (B) Barplot showing PANTHER GO Biological process categories of genes in the yellow module. (C) Barplot showing PANTHER GO Cellular Component categories of genes in the yellow module. (D) Barplot showing PANTHER Pathway categories.

**Supplementary Figure 9.** Cassava expression atlas for peroxidase family protein associated with cassava mosaic disease (CMD).

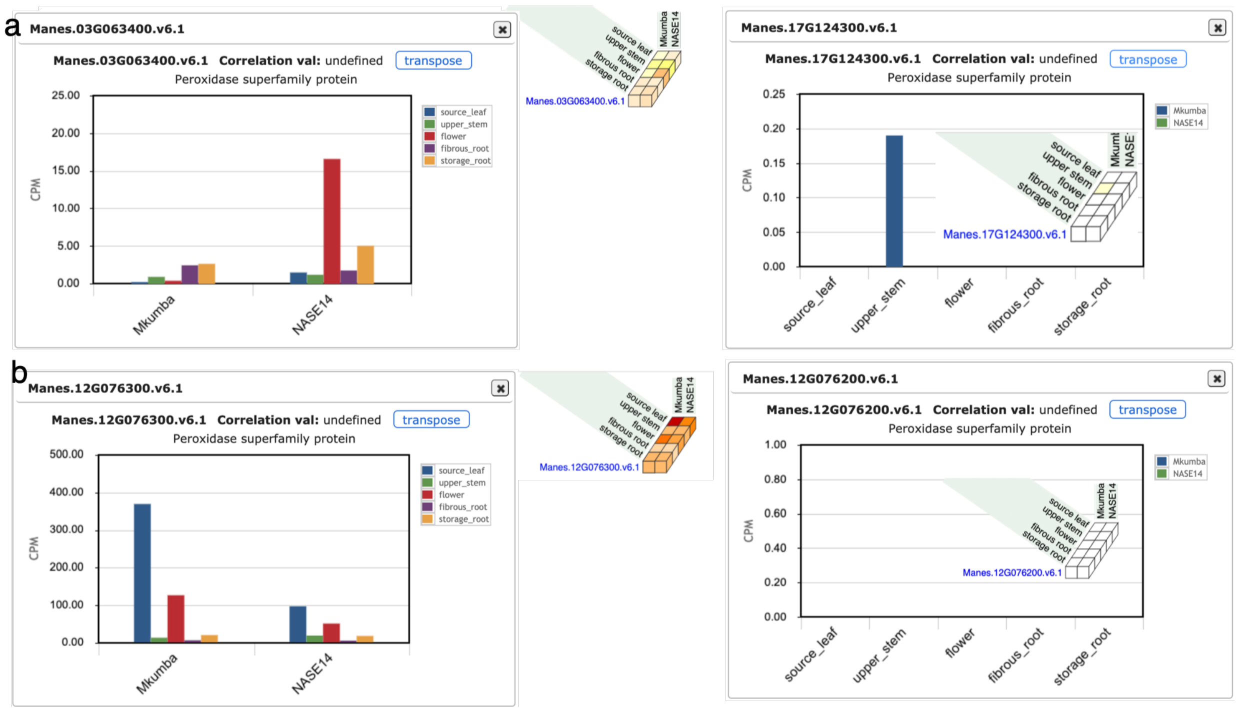


(A) The gene expression of identified peroxidase family proteins is found to be differentially expressed in the differential expression analysis of a cassava mosaic disease (CMD) resistant and susceptible cassava accessions. (B) The two peroxidase family proteins found to be associated with CMD regulation in cassava using the GWAS approach.

**Supplementary Figure 10.** Proportion of variance explained by SNPs in the DEG regions.

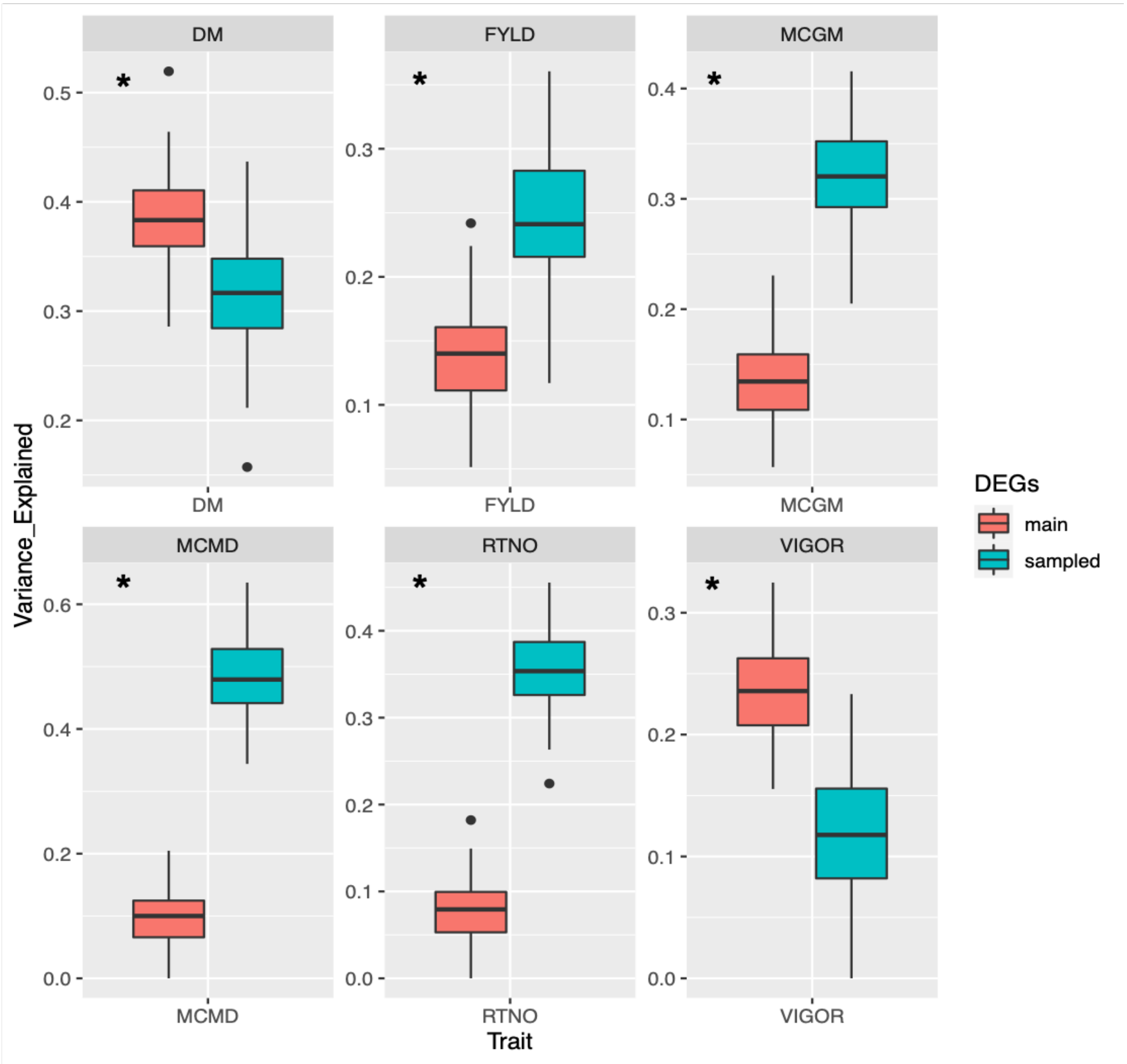


- *Significant at 0.01 by the likelihood ratio test (LRT).
- LRT = 2*(two kernel model log-likelihood – one kernel model log-likelihood)

Proportion of variance explained by the SNP markers (11,399 SNPs) in the DEG regions for agronomic traits (DM: Dry matter content, FYLD: Fresh yield, RTNO: Root number), fitness trait (VIGOR) and disease traits (MCMD: mean Severity of Cassava Mosaic Disease, MCGM: mean Severity of Cassava Green Mite). Relationship matrices were calculated using SNP markers in the DEG regions, excluding SNPs in high LD ($r2 >= 0.9$) with the SNPs in the DEG regions in the random samples of equal size and distribution. Variance components were estimated using EMMREML. The NULL model was compared to a multikernel model using a likelihood ratio test. In all cases, the multikernel models were significant at 0.01.

**Supplementary Figure 11.** Data processing and analysis schema.

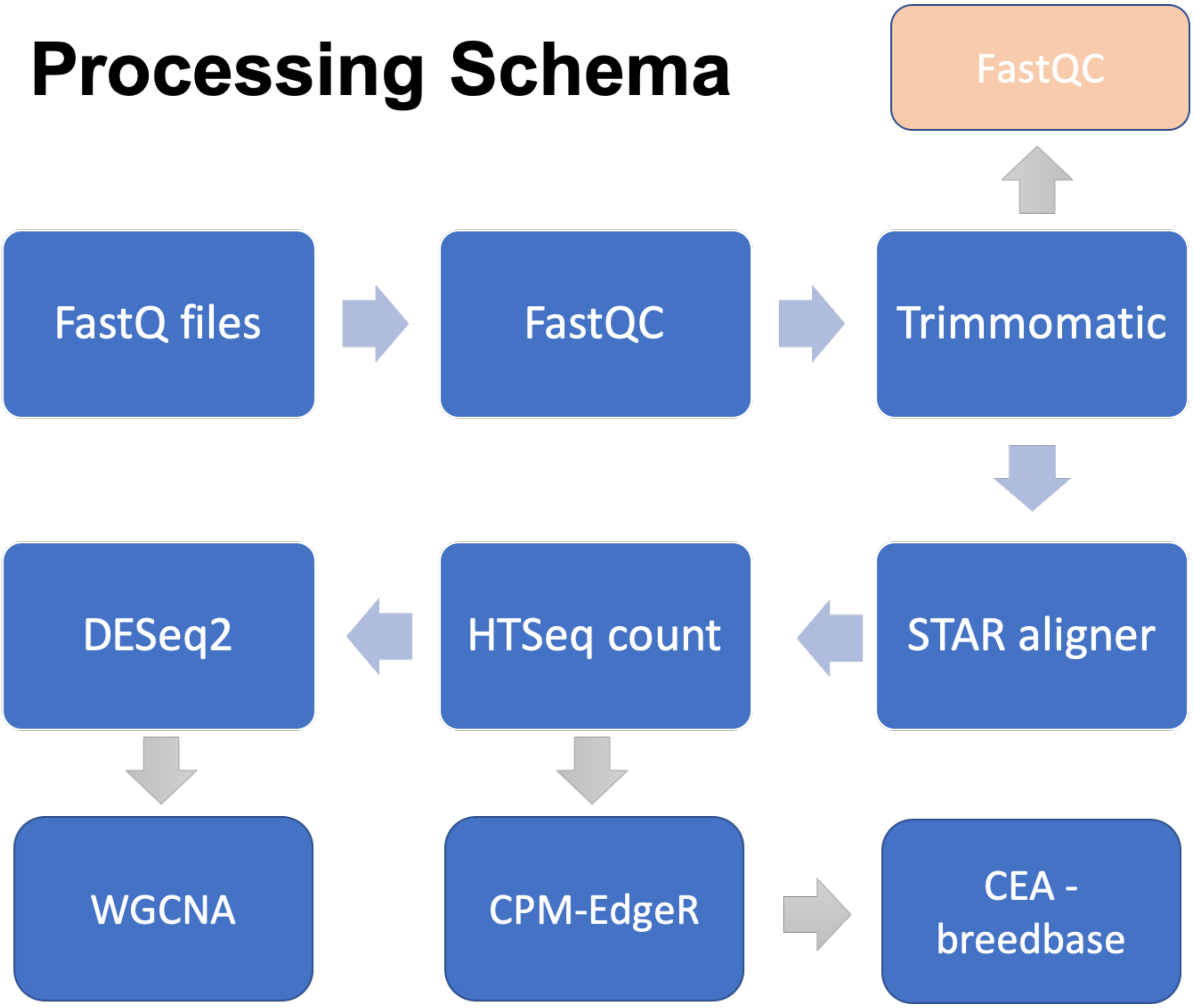


Reads generated from all cassava accessions were trimmed to remove the first 12 bp and Illumina Truseq adaptor remnants using Trimmomatic. Quality checks were done before and after trimming using FastQC. STAR aligner was used to align reads against the cassava genome annotation version 6 and gene-level counts were obtained from the resulting BAM files using HTSeq. Gene-level counts were CPM normalized using EdgeR and set up on cassava expression atlas (CEA) for gene expression visualization (https://cea.sgn.cornell.edu/expression_viewer/input). Downstream analysis was performed using DESeqs and WGCNA for differential expression analysis and weighted gene co-expression network analysis.
